# A large-scale evaluation of tree shape indices reveals potential pitfalls

**DOI:** 10.64898/2026.09.03.748783

**Authors:** Luise Häuser, Alexandros Stamatakis

## Abstract

While there exists a plethora of prior research on phylogenetic tree shape indices, a largescale analysis of the behavior of these indices on empirical trees has not yet been conducted. Here, we address this by computing 54 indices on more than 45,000,000 empirical trees retrieved from the EvoNAPS and RAxML Grove databases. To calculate the indices, we use our novel, comprehensive open-source Python library called treeshapy. The results of our large-scale evaluation indicate that there exist several potential pitfalls when conducting tree shape studies. For 14 indices we find clear indications, that they are highly sensitive to the position of the tree’s root. Only 5 indices appear stable in that sense, while we observe a medium degree of rooting instability for the remaining 35 indices under study. Therefore, uncertainties pertaining to the root placement directly affect tree shape values. Furthermore, the values of all except 5 indices are inherently correlated with tree size, even so, when applying adequate normalization techniques. As a consequence, tree shape values for trees of different sizes should generally not be compared. We further observe that several groups of indices are strongly correlated with one another. Hence, to conduct a representative tree shape study, an appropriate subset of uncorrelated indices should be selected to capture as many aspects of the tree shape as possible while avoiding essentially redundant results at the same time. Our study serves as a guide for conducting tree shape studies in a more cautious and comprehensive manner on empirical data. Apart from our open-source treehshapy Python package, we devise appropriate guidelines for selecting suitable indices and cautiously interpreting respective results.

## 1 Introduction

A plethora of indices (see Table 4) that capture and quantify phylogenetic tree shape has already been described in the literature. The distinct indices attempt to quantify different properties of tree shape. Some indices are calculated by evaluating a specific property at each tree node [17]. Others are based on the nodes depths [70, 72], on the distances between nodes [56], or on the number of nodes at the same depth [16]. Further indices rely upon the frequency of occurrence of certain characteristic subgraphs (e.g., cherries) [54]. Metrics from network science, a field related to graph theory, can also be used to quantify tree shape [12].

Tree shape indices are typically calculated in phylogenetic downstream analyses. Virologists, for instance, deploy indices to determine transmission patterns [16, 65] or to classify pathogens by their evolutionary dynamics [64, 63, 40, 5, 12]. Harcourt-Brown et al. [29] and Chindelevitch et al. [12] show, that several shape indices perform well in distinguishing trees by the type of species they contain. In addition, there exist studies that relate phylogenetic tree shapes to matrilineal fertility inheritance [9], to biodiversity skewness [31], to the interaction of plants with their pollinators [11], and to the diversification of a species after isolation [53]. Holman [35] investigate the utility of tree shape indices in the context of historical linguistics to draw conclusion about rates of language evolution. This broad range of phylogenetic downstream analysis applications illustrates the practical relevance of tree shape metrics. However, in order to correctly interpret the results, increased awareness about the mathematical properties of the indices is pivotal.

Theoretically oriented studies have extensively investigated these mathematical properties [24, 51, 7, 42]. There also exist recent comprehensive reviews of tree shape indices [24, 38]. Yet, to date, a large-scale study on the general behavior and properties of existing indices on phylogenetic trees inferred on empirical data has not yet been conducted. In most prior work, authors typically analyze but a few exemplary trees from specific application fields such as virology [31] or human genomics [9], for instance. Studies on larger tree sets have either been conducted on simulated data [38, 16] or have been limited to a small subset of specific tree shape indices [51, 7].

Therefore, we complement existing studies on tree shape indices by conducting a largescale tree shape index evaluation on trees inferred on empirical MSAs extracted from the *EvoNAPS* [67] and *RAxML Grove* [37] databases. Being based on these sources, our study covers trees for a wide variety of species. Both databases contain unrooted trees, yet most indices are defined on rooted trees only. To alleviate this, we root each tree on every possible branch. This results in a total of more than 45,000,000 rooted trees, on which we evaluate 54 implemented tree shape indices (see Fig. 1). The results of this evaluation are neither biased by a specific tree simulation model, nor by potential intrinsic properties of empirical phylogenies inferred on data from a specific field such as virology, for instance.

**Fig. 1:**
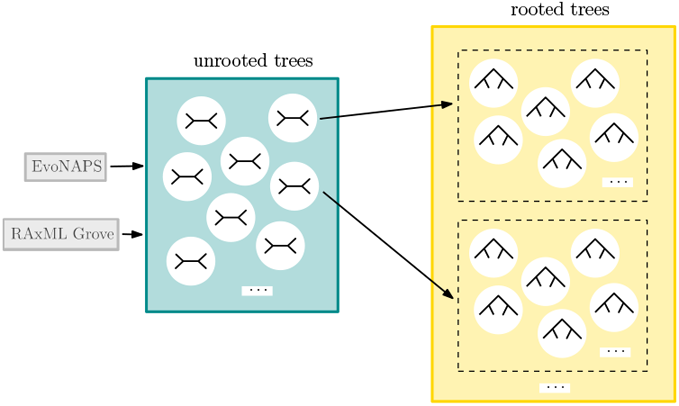
Origin of the trees in our experiments. We use almost 200,000 unrooted trees from the *EvoNAPS* [67] and *RAxML Grove* [37] databases. We root each of these empirical trees at every possible branch, resulting in more than 45,000,000 rooted trees.

Our findings provide valuable insights into index properties for which we can (currently) not devise mathematical proofs. As our study exclusively relies upon empirical data, it unveils associated issues that are of practical relevance. The results of our evaluation can hence guide and help improve the interpretation of future tree shape studies.

In particular, we investigate how the root position affects the different index values (see Section 4.6). We find that this has a pronounced effect on the values of 14 indices. 35 indices are affected in a medium degree and only for 5 indices appear to be stable with respect to the position of the root. Further, we examine the correlation between the index values and the tree size (see Section 2.3), which is again pronounced for all but 5 indices. In this context, we also describe and discuss different approaches for normalizing indices. As we show, however, these normalization techniques can only partially alleviate the observed correlation between the index values and the tree size. In Section 2.4, we further investigate the correlation of the indices with each other. The results indicate that some indices quantify tree shape in a highly analogous manner while others exploit distinct aspects of the tree shape.

Additionally, we conduct a case study (see Section 2.5) using a single phylogenetic tree inferred on DNA data for a specific receptor in different species. Via this example, we demonstrate that our general findings do apply to a specific real-world dataset. This example highlights the relevance of our findings for using tree shape indices in practice. We also introduce our open-source Python library called `treeshapy`, which we use to compute all tree shape indices.

Being the Python equivalent of the third party R library `treestats` [38], our `treeshapy` library allows for calculating 56 indices in linear time as a function of the number of leaves. Hence, it allows to seamlessly and efficiently conduct comprehensive tree shape analyses with Python.

## 2 Results

In this section, we present the results of our large-scale evaluation of 54 tree shape indices (Section 4.1) on more than 45,000,000 un-rooted trees (Section 4.5). In Section 2.1, we show our results on rooting instability. The behavior of tree shape indices defined for un-rooted trees is described in Section 2.2. Then, we elaborate on the correlation between the index values and tree size and analyze the effects of the different normalization techniques (see Section 2.3). Additionally, we examine the pairwise correlations of the indices (see Section 2.4). In Section 2.1, we present a case study illustrating the effects of rooting instabilities by means of a real-world example.

### 2.1 Rooting Instability

In the following, we investigate the rooting instability of the tree shape indices. For each index and each unrooted tree, we calculate the kurtosis of the distribution of index values across all corresponding rooted trees (see Section 4.6.1). We then analyze the distribution of the kurtosis values for each index across all unrooted trees. Fig. 2 hence contains a box plot for each index. The indices are classified as described in Section 4.1, and the box plots are colored accordingly. Note that the y-axis is logarithmically scaled. One dashed line at level 1.8 indicates the kurtosis of the uniform distribution, a second dashed line at level 3, denotes the kurtosis of the normal distribution. If the kurtosis values of an index are below 1.8, this suggests that the index is stable under distinct rootings. A kurtosis between 1.8 and 3 points to some instabilities, while values above 3 indicate substantial rooting instability. In additional analyses we subdivide the rooted trees according to whether the root is placed on an external or internal branch (see Fig. 17 and Fig. 16 in the appendix). This helps to more precisely analyze the behavior depending on the root position.

**Fig. 2:**
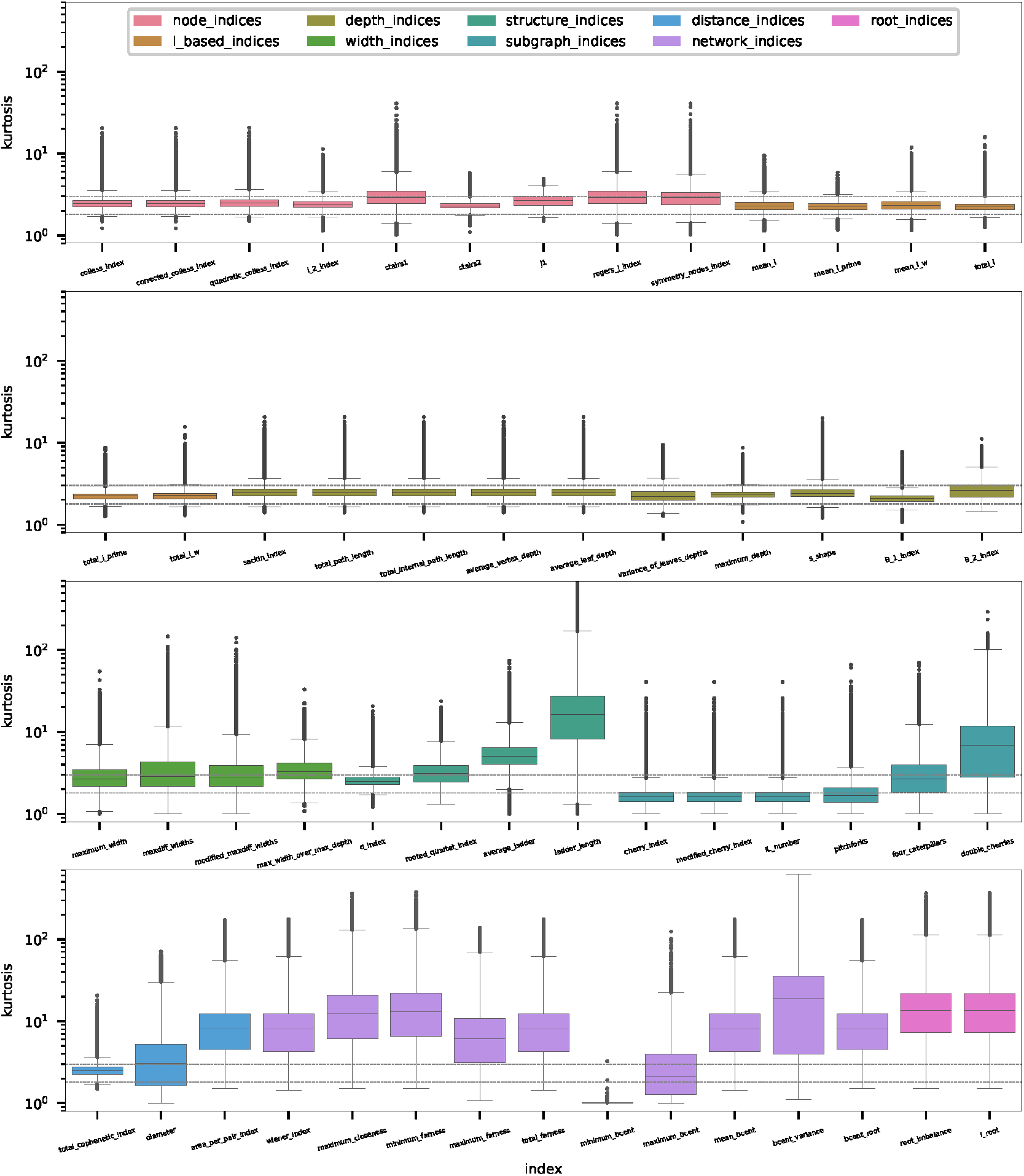
Kurtosis distributions for all tree shape indices over the trees under study. The plots are grouped and colored according to the underlying approach of the indices (see Section 4.1). A dashed line at level 1.8 (3) indicates the kurtosis of the uniform (normal) distribution. The y-axis is logarithmically scaled.

Our experiment shows that the indices under study exhibit varying degrees of rooting instability. In the following, we describe and discuss the main observations for the different index categories (see Table 4). To develop a better intuition for our findings, we also examine how moving the root from one branch to an adjacent one affects the values of the various indices.

We observe medium kurtoses (mostly between 1.8 and 3) for all node indices and *I*-based indices. These indices are summary statistics of properties that are calculated individually for each internal node. The node properties examined consider only the subtree below the respective node. If we move the root from one branch to an adjacent branch, this only affects the subtree below a single node *v* (see Fig. 3). Consequently, moving the root only affects the node properties of *v*, while the properties of all other nodes remain unchanged. The resulting index value distributions therefore exhibit some but not a pronounced variation. The corresponding kurtosis values indicate medium rooting instability for node indices and *I*-based indices. The kurtoses tend to be slightly higher for discrete indices (e.g., `rogers_j_index`) than for continuous indices (e.g., `colless_index`).

**Fig. 3:**
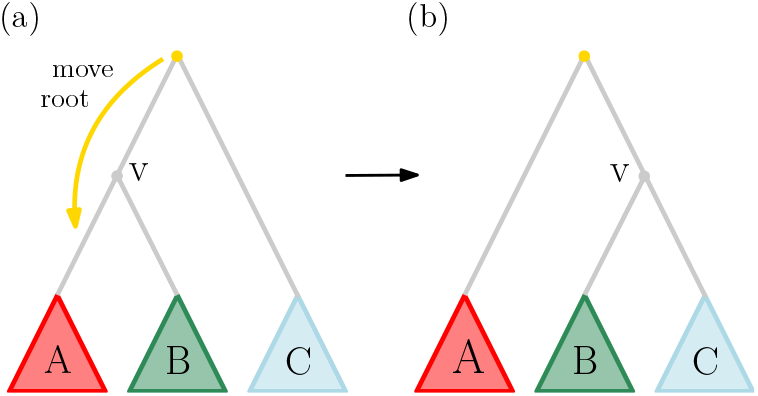
Moving the root from a branch to a neighboring branch. Moving the root only affects the node properties of *v*, while the properties of all other nodes remain unaltered. It increases the depth of all nodes in subtree *C* by 1, while decreasing the depth of all nodes in *A* by 1. The depth of *v* and the nodes in *B* remains unchanged.

The kurtosis is also at a medium level for depth indices. Roughly, calculating a depth index on a tree is equivalent to computing a specific summary statistic over the depths of its nodes. Moving the root from one branch to another increases the depth of the nodes in one subtree, while decreasing the depth of the nodes in another subtree. For a third group of nodes, the depth remains unchanged (see Fig. 3). The depth indices are therefore only slightly affected by moving the root, particularly so, when the affected subtrees are of similar size. Consequently, the depth indices predominantly exhibit kurtosis values below 3, indicating a moderate degree of rooting instability. The differences between the various depth indices can be explained by the different summary statistics used to calculate each index.

The kurtosis distributions of the width indices exhibit more substantial variance. The medians are close to or slightly above 3, suggesting increased rooting instability. The width indices are related to the depth indices, since we determine the widths by grouping the nodes according to their depth. Although moving the root to a neighboring branch can change the depth of a single node by at most 1, the effect on the width can be more pronounced. Fig. 4 illustrates an example in which the `maximum_width` is decreased by 4 by moving the root from one branch to a neighboring branch. It is intuitive, that even larger differences can occur in larger trees. On the other hand, Fig. 5 shows that the `maximum_width` must not necessarily be affected by moving the root. This explains, why the kurtosis distributions of the width indices do not only exhibit higher medians but also an increased variance.

**Fig. 4:**
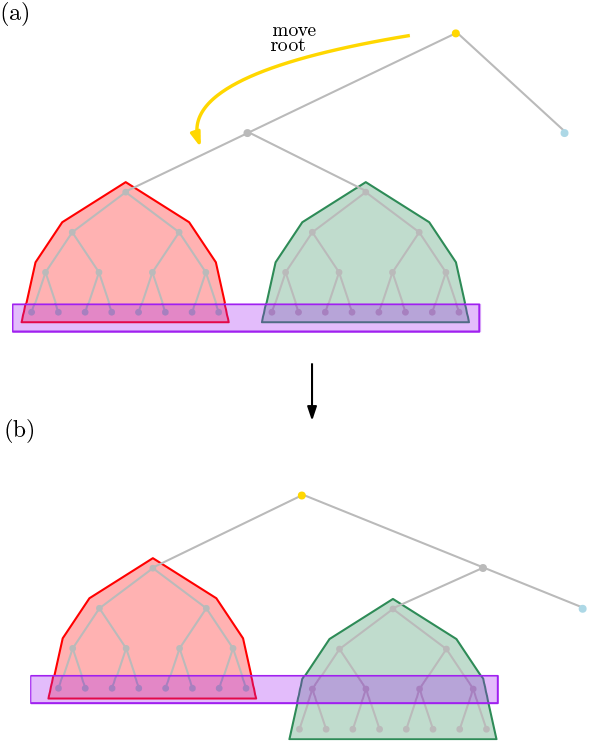
Impact on maximum_width when moving the root from a branch to a neighboring branch (1). Subtrees are colored according to Fig. 3.`maximum_width` decreases from 16 (a) to 12 (b). Corresponding nodes are highlighted in purple.

**Fig. 5:**
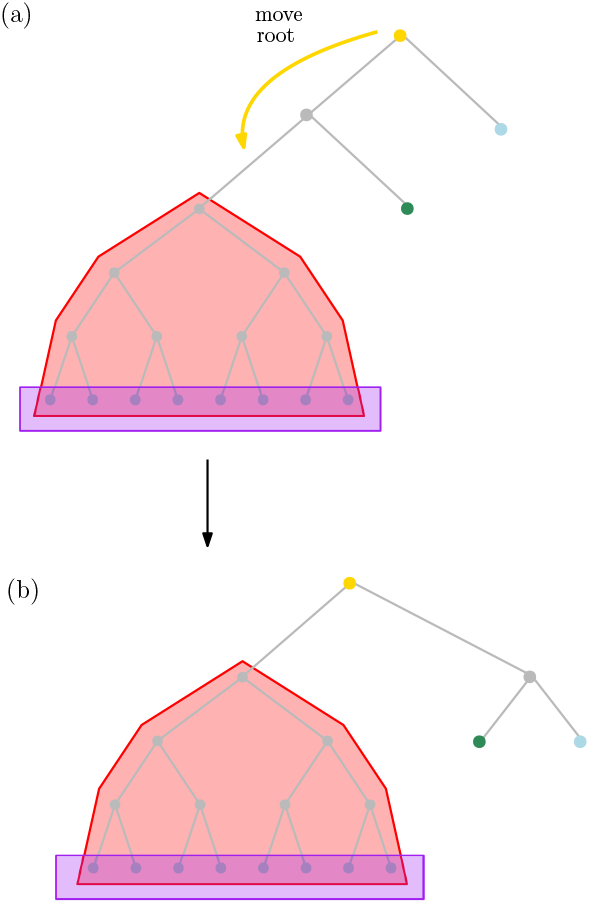
Impact on the maximum_width when moving the root from a branch to a neighboring branch (1). Subtrees are colored according to Fig. 3. `maximum_width` is 12, both in (a) and in (b). Corresponding nodes are highlighted in purple.

The structure indices behave differently when compared to each other. `d_index` yields medium kurtosis values, just as the depth indices. `rooted_quartet_index` appears more unstable, comparable to the width indices. For `average_ladder`, the values are even higher. We focus on the `ladder_length` for which we observe exceptionally high kurtosis values. We explain this behavior via the following example: Let ℛ (*T* ) be the set of rooted trees corresponding to the unrooted tree *T* . Let *m* be the maximum `ladder_length` among all trees in *R*(*T* ) (see Fig. 6 (a)). We refer to a ladder of length *m* as the maximum ladder. We further assume that the maximum ladder is the only ladder of length *>* 1. For the rooted trees in ℛ (*T* ), we can distinguish between two cases: In the first case, the branch on which the root is placed does not form part of the maximum ladder. The maximum ladder is thus contained in this rooted tree, which therefore exhibits `ladder_length` *m*. In the second case, the tree is rooted at a branch that forms part of the maximum ladder. Consequently, the maximum ladder breaks (see Fig. 6 (b)). The `ladder_length` of the rooted tree then corresponds to the length of the larger part of the maximum ladder that is preserved. This leads to a distribution of index values which has a clear peak at *m* but a tail diminishing toward 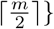, which induces a high kurtosis.

**Fig. 6:**
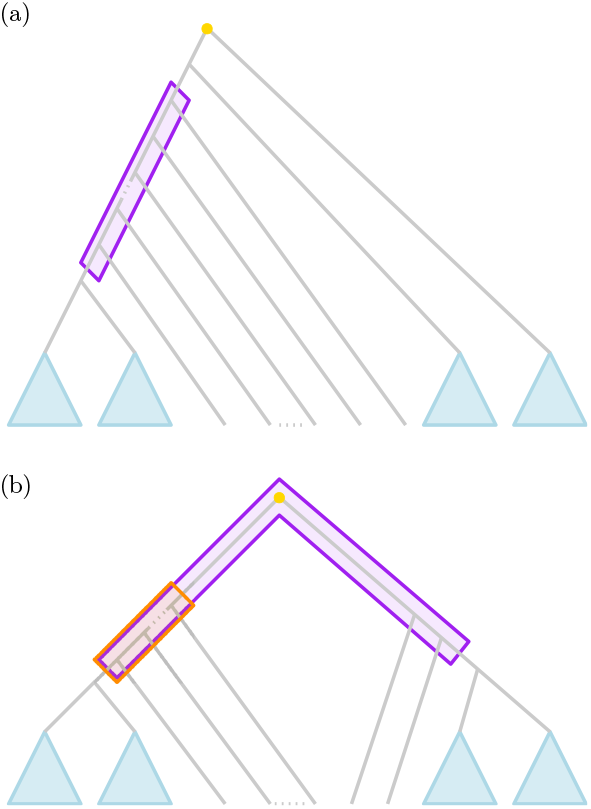
Effect of the root’s position on ladder_length. (a): Rooted tree with maximum ladder (highlighted in purple), blue subtrees are fully balanced binary trees only containing trivial ladders (b): Topology from (a) with the root placed on a different branch such that the maximum ladder from (a) breaks. The maximum ladder in this topology is highlighted in orange.

For subgraph indices, it can be observed that kurtosis increases with the size of the relevant subgraph (see Fig. 7): It is mostly below 1.8 for `cherry_index`, `modified_cherry_index`, and `il_number`, but substantially higher for `double_cherries`. A subgraph index is based on counting the number of occurrences of the respective subgraph in the tree. Moving the root changes the value of such an index by 1 if it either breaks or creates such a subgraph (see Fig. 8). The larger the relevant subgraph, the less frequently it tends to occur. When the corresponding index changes by 1, the relative impact is higher. Consequently, the subgraph indices associated with larger subgraphs yield higher kurtoses and appear to be less stable with respect to the root position. Note also that the number of cherries remains constant as long as the tree is rooted on an internal branch. Consequently, `cherry_index`, `modified_cherry_index`, and `IL_number` are completely stable for internal roots (see Fig. 16).

**Fig. 7:**
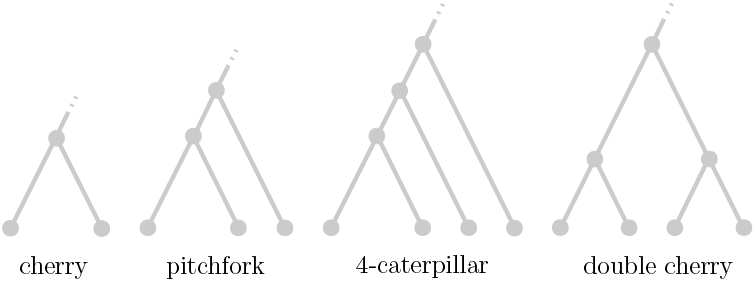
Subgraphs counted to evaluate subgraph indices. `IL_number` and `modified_cherry_index` also rely on the number of cherries.

**Fig. 8:**
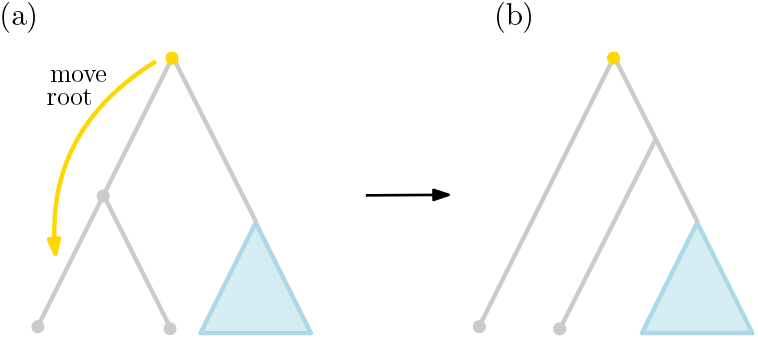
Effect of moving the root on the. `cherry_index`. In (a), there is a cherry which does not exist in (b) due to the altered position of the root. Note that the number of cherries is not affected as long as the root is placed on an internal branch.

The `total_cophenetic_index` is a distance index, but behaves analogously to the depth indices with respect to kurtosis. The index is based on the pairwise discrete patristic distances among the leaves. Such a distance depends on which node is the least common ancestor of the leaves under consideration. Referring to the example in Fig. 3, moving the root only affects the patristic distance of leaf pairs in which one leaf is in subtree *B* and the other is in subtree *A* or *C*. Moving the root therefore only slightly changes the value of the `total_cophenetic_index`, in analogy to the depth indices. This leads to kurtosis values that mostly range between 1.8 and 3, indicating moderate rooting instability. In contrast, both the `diameter`, and the `area_per_pair_index`, yield higher kurtoses. Since these indices are based on distances across the entire tree, this behavior can be explained in the same way as we do below for the network indices.

Also, for most network indices, we observe high kurtoses. When calculating such a network index, we evaluate a property from network science for each node individually and obtain the index value using a summary statistic. The approach is mostly analogous to the computation of node indices. However, in this case, the tree is considered to be a network. Consequently, when calculating a node’s property, the entire tree is taken into account, not just the subtree below that node. Furthermore, considering the tree as a network means that the root is treated like an ordinary internal node. Therefore, a shift in the root’s position changes the network properties of numerous nodes. In particular, this affects all cases where the shortest path between two nodes traverses the root. For most network indices, this results in high kurtosis. Only the `minimum_bcent` and `maximum_bcent` indices exhibit low and medium kurtoses, respectively. These indices are based on a property called *betweenness centrality*, which corresponds to the number of shortest paths passing through the respective node [12]. Shifting the root can affect the betweenness centrality of individual nodes, while the overall minimum and maximum remain unaltered, resulting in distributions with the observed, comparatively low kurtosis values.

The kurtosis values observed for root indices are high. These indices solely capture the balance of the root node itself. This means that the calculation only takes into account the *number* of leaves present in the two subtrees below the root’s immediate children. All other properties of these subtrees are ignored. Hence, intuitively, root indices generally exhibit high kurtosis values. However, if the root is located on an external branch, the smaller subtree always comprises but a single leaf, while the larger one encompasses all remaining leaves. Consequently, root indices for external roots are entirely stable (see Fig. 17).

Overall, this study points to moderate rooting instabilities for the vast majority of examined tree shape indices. Only for a few indices (`cherry_index`, `modified_cherry_index`, `IL_number`, `pitchforks`, and `minimum_bcent`), we observed a low instability. High kurtoses occur for `average_ladder`, `ladder_length`, `double_cherries`, `area_per_pair_index`, almost all network indices as well as root indices, indicating a high degree of rooting instability.

While such a high kurtosis is a clear indicator of instability, low kurtosis does not necessarily mean that the index is completely stable. If the value distribution of an index has a kurtosis of 1.8, this could mean that the values are uniformly distributed across all possible rooted versions of an unrooted tree. Intuitively, one would interpret this as low rooting instability as long as the range of the uniform distribution is sufficiently small. However, kurtosis is independent of the nominal values and does therefore not allow to draw conclusions about this range. Consequently, the range of the uniform distribution can theoretically be arbitrarily large, which means that the rooted versions of an unrooted tree can have uniformly distributed, yet highly different index values. Such an index does therefore not behave in a way that we would describe as stable, despite the fact that it exhibits a comparatively low kurtosis. Our case study in Section 2.5 provides some insights in that direction.

### 2.2 Indices defined for Unrooted Trees

In an additional study, we focus on the behavior of the indices defined on both rooted and unrooted trees. We not only evaluate these indices on all rooted versions of each unrooted tree, but also on the original unrooted tree itself. We determine the percentile rank of the resulting index value for the unrooted tree relative to the index values obtained for all rooted versions of that tree. If the percentile rank is 0, the value for the unrooted tree is lower than all index values obtained for the corresponding rooted trees. Conversely, the index value for the unrooted tree is highest when the percentile rank is 100. For each index, we determine the mean and standard deviation of these percentile ranks across all unrooted trees examined (see Table 1).

**Tab. 1.** Percentiles for unrooted tree relative all rooted versions of that tree.

| index | mean | std. dev. |
| --- | --- | --- |
| diameter | 36.342 | 9.365 |
| area_per_pair_index | 0.0 | 0.0 |
| wiener_index | 0.0 | 0.0 |
| maximum_closeness | 99.979 | 1.448 |
| minimum_farness | 0.0 | 0.0 |
| maximum_farness | 0.0 | 0.0 |
| total_farness | 0.0 | 0.0 |
| minimum_bcent | 52.212 | 3.874 |
| maximum_bcent | 0.0 | 0.0 |
| mean_bcent | 9.262 | 6.843 |
| bcent_variance | 0.069 | 0.83 |

This table refers to the percentile ranks of the index value for the unrooted tree relative to the index values obtained for all rooted versions of that tree. It shows mean and standard deviation of these percentile ranks over all un-rooted trees under study.

For 6 indices (`area_per_pair_index`, `wiener_index`, `maximum_bcent`, and all indices based on farness), the percentile rank is always 0, that is, the value resulting for the rooted tree is always the lowest. For `maximum_closeness`, it is alwayst the highest, apart from a few large trees in which numerical instabilities occur (percentile rank close to 100). For the remaining 4 indices (`diameter`, `minimum_bcent`, `mean_bcent`, `bcent_variance`), the behavior is less clear.

The *farness* of a node is defined as the sum over the discrete distances to all other nodes. If the tree is rooted, the farness of a node *v* increases by 1 (compared to the unrooted tree) for every node *w* for which the shortest path from *v* to *w* passes through the root. Rooting thus increases the farness by at least 1 for every node. Consequently, all metrics based on farness are higher for rooted trees than for the original unrooted tree. The same argument applies to the `area_per_pair_index` and the `wiener_index`, if one follows the definitions of these indices. *Closeness* is defined as the reciprocal of farness. Consequently, the closeness of each node in a rooted tree is lower than in the corresponding unrooted tree. Therefore, the `maximum_closeness` is also lower for rooted trees.

The *betweenness centrality* (bcent) of a node *v* corresponds to the number of shortest paths that pass through *v*. Adding a root to an unrooted tree changes the betweenness centrality of *v*, if the shortest path from the root to another node *w* passes through *v*. This does not necessarily apply to all nodes, but it always affects the node with the highest betweenness centrality. Therefore, the `maximum_bcent` is higher for any tree with a specified root than for the unrooted tree. For the other indices based on betweenness centrality, the behavior is less clear. The `diameter` is also affected only if the longest shortest path in the tree passes through the root. As a result, different percentile ranks occur for this index as well.

Overall, this analysis of unrooted indices shows, that the index values resulting for un-rooted trees cannot be compared directly to the results obtained for (corresponding) rooted trees. However, these indices defined for un-rooted trees all exhibit high rooting instabilities (see Section 2.1) when being evaluated on rooted trees. Against this background, it is preferable to evaluate them directly on the unrooted tree.

### 2.3 Correlation with Tree Size

In general, the absolute values of a tree shape index cannot be compared across trees of different sizes, as they may depend on the number of tips in the tree. Normalization aims to eliminate (or at least reduce) this dependency and yield the values comparable. To this end, we have introduced three approaches (see Section 4.3): normalization by maximum, normalization by the expected value under the Yule model, and normalization by number of tips.

Janzen and Etienne [38] conduct a study on simulated data to quantify how well the different normalization techniques eliminate tree size dependency. They use four distinct diversification models that differ in their evolutionary rates. They simulate 10,000 replicates per model and tree size. They find that across all simulation models, normalization by number of tips is successful for `pitchforks` and normalization by expected value under the Yule model is successful for `cherry_index` and `rooted_quartet_index`. The latter normalization technique also performs well for `area_per_pair_index`, `sackin_index`, `colless_index`, and `total_cophenetic_index`, but only for data simulated under the Yule model. In all other cases, they do not observe a successful normalization. However, they limit their study to but a few indices and to only one normalization technique per index. Furthermore, they do not use empirical tree data, which may behave differently from simulated data. We therefore use the results of our large-scale evaluation to investigate the effectiveness of the different normalization techniques under a complementary setting.

For each unrooted tree *T* under study, we create a set ℛ (*T* ) of rooted trees by rooting *T* at every possible position (see Section 4.6). For all rooted trees constructed in this manner, we do not only determine the absolute value of each index, but also the normalized values according to the three different normalization techniques (whenever the necessary properties of the respective index for computing the normalization are available). For each index, we then determine the Spearman rank correlation of the absolute and of the different normalized values with the tree size. Thereby, we investigate, to which extent the index values depend on the input tree size. If there is a clear dependency, the index values cannot be compared for trees of different size. We decided to focus on rank correlation because tree shape indices are often used to compare different topologies, while the exact numerical values are disregarded. For the sake of completeness, we provide the results using the Pearson correlation in the appendix (see Table 9).

The results using Spearman rank correlation are provided in Table 2. To improve readability, the cells are colored based on their values. Cells with a correlation *<* 0.1 are colored in green. The low correlation indicates that there is almost no tree size dependency; normalization is hence either not required, or not successful. Cells with values between 0.1 and 0.3 are colored in yellow, those with values between 0.3 and 0.6 in orange. Red cells contain values *>* 0.6 indicating a strong dependency on tree size. We observe that, for almost all indices, the absolute values strongly correlate with tree size. Only for 5 indices (`stairs1`, `j1`, `B_2_index`, `ladder_length`, and `I_root`), the absolute values exhibit a correlation with tree size which is not greater than 0.3.

**Tab. 2.** Spearman rank correlation of index values with tree size.

| index | abs. | rel.(tips) | rel.(max) | rel.(yule) |
| --- | --- | --- | --- | --- |
| colless_index | 0.82 | 0.42 | -0.62 | 0.33 |
| corrected_colless_index | -0.64 | -0.86 | -0.62 | -0.7 |
| quadratic_colless_index | 0.83 | 0.7 | -0.46 | 0.65 |
| I_2_index | -0.41 | -0.94 |  |  |
| stairs1 | -0.3 | -0.96 | -0.3 |  |
| stairs2 | 0.38 | -0.93 |  |  |
| j1 | -0.25 | -0.86 |  |  |
| rogers_j_index | 0.94 | -0.23 | -0.3 |  |
| symmetry_nodes_index | 0.95 | -0.21 | -0.28 |  |
| mean_I | -0.39 | -0.94 |  |  |
| mean_I_prime | -0.39 | -0.94 |  |  |
| mean_I_w | -0.38 | -0.93 |  |  |
| total_I | 0.89 | -0.36 |  |  |
| total_I_prime | 0.88 | -0.37 |  |  |
| total_I_w | 0.87 | -0.36 |  |  |
| sackin_index | 0.88 | 0.52 | -0.55 | 0.35 |
| total_path_length | 0.88 | 0.52 | -0.55 |  |
| total_internal_path_length | 0.87 | 0.52 | -0.55 |  |
| average_vertex_depth | 0.52 | -0.77 | -0.55 |  |
| average_leaf_depth | 0.52 | -0.79 | -0.55 | -0.34 |
| variance_of_leaves_depths | 0.31 | -0.43 |  | -0.28 |
| maximum_depth | 0.51 | -0.81 | -0.7 |  |
| s_shape | 0.93 | -0.06 |  |  |
| B_1_index | 0.96 | 0.45 |  |  |
| B_2_index | 0.05 | -0.78 | -0.09 | 0.5 |
| maximum_width | 0.82 | -0.36 |  |  |
| maxdiff_widths | 0.64 | -0.42 |  |  |
| modified_maxdiff_widths | 0.65 | -0.45 |  |  |
| max_width_over_max_depth | 0.49 | -0.47 |  |  |
| d_index | 0.53 | -0.73 |  |  |
| rooted_quartet_index | 0.93 | 0.91 |  | -0.79 |
| average_ladder | -0.39 | -0.9 |  |  |
| ladder_length | -0.03 | -0.8 |  |  |
| cherry_index | 0.94 | 0.39 | 0.43 | 0.39 |
| modified_cherry_index | 0.89 | -0.39 | -0.43 |  |
| IL_number | 0.89 | -0.39 |  |  |
| pitchforks | 0.89 | 0.06 |  |  |
| four_caterpillars | 0.85 | 0.06 |  |  |
| double_cherries | 0.65 | 0.09 |  |  |
| total_cophenetic_index | 0.86 | 0.74 | -0.46 | 0.65 |
| diameter | 0.67 | -0.88 |  |  |
| area_per_pair_index | 0.82 | -0.94 |  | -0.16 |
| wiener_index | 0.98 | 0.96 |  |  |
| maximum_closeness | -0.93 | -0.97 |  |  |
| minimum_farness | 0.93 | 0.76 |  |  |
| maximum_farness | 0.93 | 0.76 |  |  |
| total_farness | 0.98 | 0.96 |  |  |
| minimum_bcent | 0.67 | 0.51 |  |  |
| maximum_bcent | 0.99 | 0.96 |  |  |
| mean_bcent | 0.95 | 0.82 |  |  |
| bcent_variance | 0.97 | 0.96 |  |  |
| bcent_root | 0.43 | 0.31 |  |  |
| root_imbalance | 0.45 | -0.96 | 0.08 |  |
| I_root | 0.08 | -0.89 | 0.08 |  |

The cells are colored according to their values ( $< 0.1$ : green, $[0.1, 0.3]$ : yellow, $[0.3, 0.6]$ : orange, $> 0.6$ : red). If a cell is gray, the normalization technique is not applicable for the respective index (see Section 4.3).

Normalizing the index value by the number of tips helps reduce the correlation for some indices (e.g., `rogers_j_index`, `symmetry_nodes_index`, and the subgraph indices based on larger subgraphs). However, there are also cases in which the normalized index exhibits a higher correlation with tree size than the absolute index (e.g., mean I-based indices, many node indices). We advise against using this normalization approach, especially if a different normalization is applicable for the respective index.

Normalization using the maximum value or the expected value under the Yule model reduces the correlation with the size for the vast majority of indices for which it is defined. Despite this positive effect, there exist indices (such as the `rooted_quartet_index`) for which the absolute values and all normalized values examined heavily depend on tree size. Please note that our experiments only measure monotonic correlations. If an index correlates with tree size in any other manner (e.g. based on axis symmetries), this may not be captured in our analysis. Additionally, we confirm the indices correlation with tree shape via a principle component analysis (PCA).

When comparing index values between trees of different sizes, we therefore strongly recommend to check that the correlation between these values and the tree size is sufficiently low. This also applies to values that have been normalized using one of the three normalization techniques we considered. If a significant correlation is detected, a comparison is not feasible.

### 2.4 Correlation of Indices

We also investigate which tree shape indices are closely related to each other. A similar behavior suggests that the corresponding indices quantify the same tree shape property. For each index *I*, we consider *I*_all_ := ⋃_*T* ∈database_ *I*(*T* ) and determine the pairwise absolute Spearman rank correlations of these value sets for all indices under study. We again focus on rank correlation, since in most experiments, the index values are compared for different topologies, while the numerical values are only of minor importance. In the resulting plot (see Fig. 9), the indices with high correlations are grouped together. A plot in which the indices are ordered based on the conceptual approach can be found in the appendix (see Fig. 19).

**Fig. 9:**
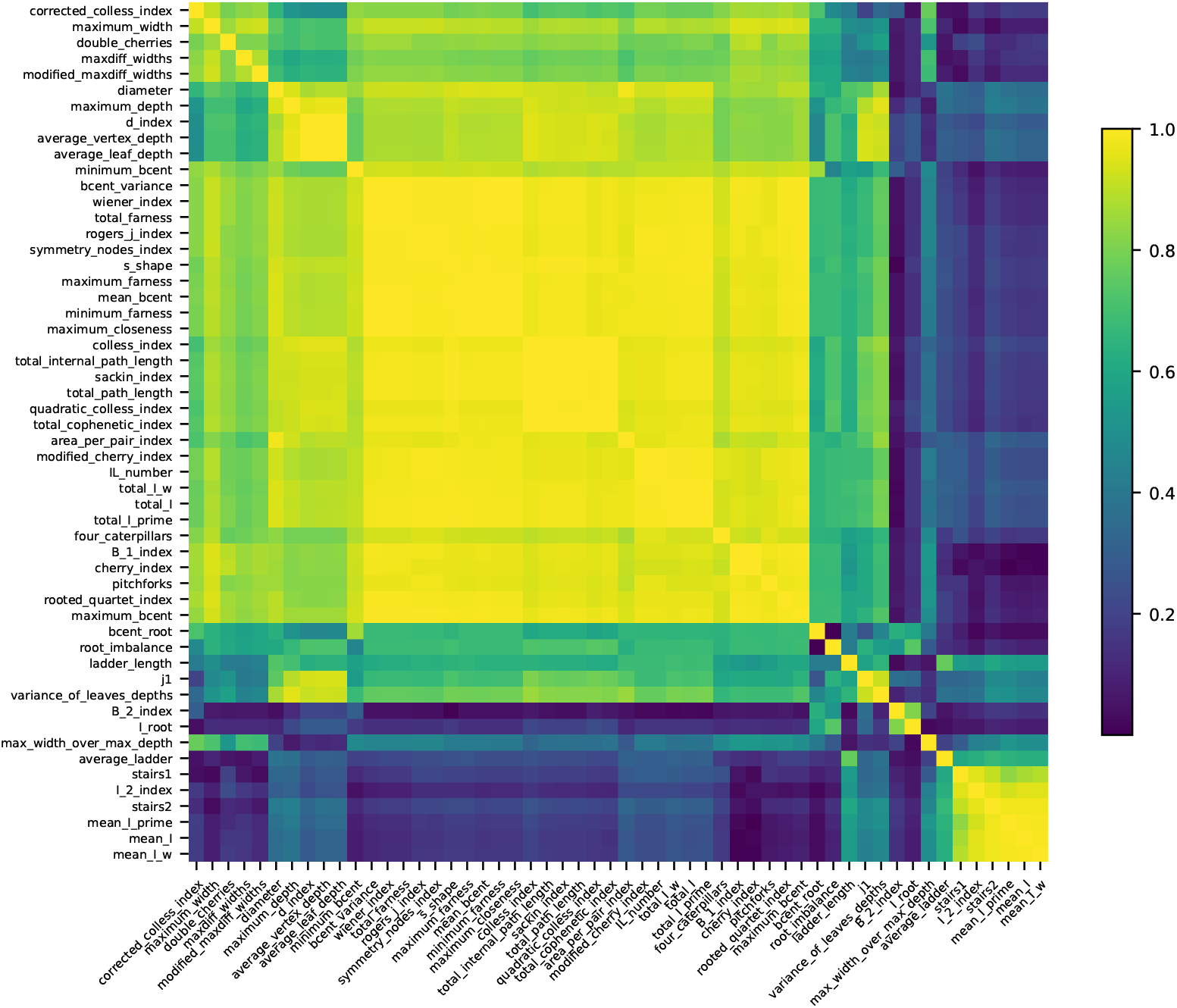
Pairwise Spearman Rank correlations of tree shape indices. Indices with high correlations are grouped together. The correlations are obtained using all rooted trees under study.

We observe groups of indices that exhibit substantial pairwise correlations. The first group comprises nearly half of the indices under study (`bcent_variance` through `maximum_bcent`). These include numerous node indices, network indices, and subgraph indices, as well as all distance indices and those I-based indices that compute a total value. In particular, we observe a high correlation between the `colless_index` and the `sackin_index`, which is consistent with the results of Lemant et al. [49]. The `quadratic_colless_index` and the `total_cophenetic_index` also exhibit a high correlation, as previously observed by Kersting et al. [43]. Additional indices show a moderate correlation with the indices in the first group (`corrected_colless_index` through `minimum_bcent`). The second group (`stairs1` to `mean_I_w`) of highly correlated indices contains some node indices and the I-based indices, which correspond to a mean value. Apart from these groups, there are indices that do not exhibit a significant correlation with other indices, such as `ladder_length` or `bcent_root`.

Overall, there are numerous index pairs with high correlations (*>* 0.9). This implies that evaluating both of them will yield redundant results. However, indices relying on the same approach are not always correlated. Inversely, there also exist indices which are heavily correlated, despite relying on distinct concepts (see also Fig. 19). We recommend selecting a subset of indices with low pairwise correlations. Thereby, distinct tree shape characteristics can be captured and analyzed via a small and manageable number of indices. To this end, for *k* ∈ [2, 10], we determine index sets of size *k* such that the selected indices exhibit minimal pairwise rank correlations in our experiments (see Appendix D.1 in the appendix).

### 2.5 Case Study

To investigate rooting instability in a real-world example, we conduct a case study on an exemplary (unrooted) tree, which we call the receptor tree. In the receptor tree, there exist two branches for which the probability, that the root is located on that branch, is not negligible. We denote the trees rooted on these branches by *T*_1_ and *T*_2_ (see Section 4.7 for details).

Many practical applications of tree shape indices compare the results of different trees [29, 31, 65], since stand-alone tree shape values are difficult to interpret. We hence use the simple and widely used Yule model to sample 10,000 unrooted trees that have the same size as the receptor tree (24). We root each of these trees on every possible branch and again calculate the tree shape indices for all resulting rooted trees (see Fig. 10). Note that we limit ourselves to trees of size 24, as most indices correlate with tree size and normalization is challenging (see Section 4.3). The 10,000 sampled trees only cover a tiny fraction of the tree space which comprises ∼2.53 × 10^28^ unrooted trees for 24 taxa. However, this number suffices to outline the practical implications of our findings on the behavior of tree shape indices.

**Fig. 10:**
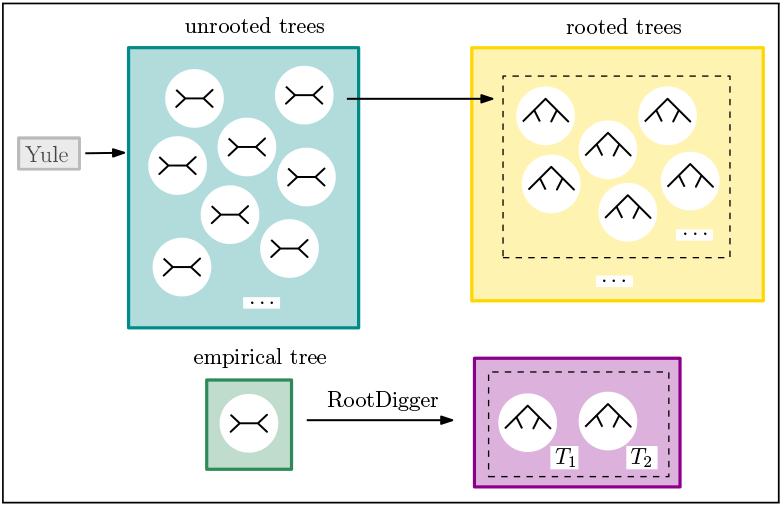
Experimental setup of our case study.

For each index, we determine the percentile ranks of the rooted receptor tree (*T*_1_, *T*_2_, respectively) with respect to the rooted trees we sampled via the Yule model. If a rooted receptor tree yields a percentile rank of 0, its index value will be lower than the index values of all sampled rooted trees. Inversely, if we obtain a percentile rank of 100, the index value the rooted receptor tree is higher than for all other observed index values. If the percentile ranks of the two differently rooted receptor trees (*T*_1_ and *T*_2_) differ substantially, this indicates a rooting instability of the respective index.

As a possible approach to handle rooting instability, we examine the mean index values of all rooted trees that correspond to the unrooted receptor tree. We consider both, the unweighted mean, and the mean weighted by the LWR of the respective root position. In the case of the receptor tree, this weighted mean is very close to the weighted mean of the values of *T*_1_, and *T*_2_, since the LWRs of all other putative rooting positions (root placement branches) are negligible. For each index we determine the percentile ranks of the unweighted and the weighted mean as described above.

In Table 3, we present the resulting percentile ranks for *T*_1_, and *T*_2_. We observe a slight trend for the rooted receptor trees to yield high percentile ranks for imbalance indices and low percentile ranks for balance indices (see Table 6). This means that the receptor tree is assessed to be less balanced than the sampled trees. This is in line with other studies where the authors also find, that trees sampled via the Yule model tend to be more balanced than empirical ones [9, 32, 27, 28].

**Tab. 3.** Results of the case study on a tree inferred for a specific receptor gene. *T*_1_, and *T*_2_ result from rooting the unrooted receptor tree at the two by far most likely positions with an accumulated likelihood weight of 0.933. For each index and each of the two trees *T*_1_ and *T*_2_, we calculate the percentile ranks relative to 10,000 trees sampled with the Yule model (see Section 4.7). Additionally, we calculate the unweighted and the LWR-weighted mean for each index over all possible rootings of the unrooted receptor tree and determine the percentile ranks with the same method. The percentile ranks of *T*_1_, and *T*_2_ are highlighted in bold if they differ by more than 10 from each other, which indicates a substantial impact of root placement on tree shape analysis results for our specific example.

| index | $T_1$<br>(0.780) | $T_2$<br>(0.153) | mean<br>(unw.) | mean<br>(LWR-w.) |
| --- | --- | --- | --- | --- |
| colless_index | **60.8** | **35.1** | 66.0 | 75.6 |
| corrected_colless_index | **72.5** | **44.7** | 76.6 | 85.2 |
| quadratic_colless_index | **46.6** | **22.3** | 50.5 | 61.5 |
| I_2_index | 81.9 | 80.6 | 85.8 | 85.7 |
| stairs1 | 77.3 | 77.3 | 92.6 | 77.3 |
| stairs2 | 11.0 | 19.0 | 10.0 | 7.4 |
| j1 | **37.6** | **66.3** | 36.3 | 24.7 |
| rogers_j_index | 44.1 | 44.1 | 77.3 | 77.3 |
| symmetry_nodes_index | 41.2 | 41.2 | 75.9 | 75.9 |
| mean_I | 88.5 | 82.3 | 86.4 | 90.2 |
| mean_I_prime | 87.9 | 79.7 | 86.1 | 90.7 |
| mean_I_w | 91.5 | 86.8 | 89.7 | 92.6 |
| total_I | **82.4** | **65.9** | 77.4 | 85.0 |
| total_I_prime | **81.7** | **63.8** | 77.1 | 85.2 |
| total_I_w | **86.1** | **72.1** | 82.0 | 88.1 |
| sackin_index | **47.7** | **22.7** | 55.4 | 65.1 |
| total_path_length | **49.1** | **23.8** | 56.8 | 66.4 |
| total_internal_path_length | **50.5** | **24.9** | 58.2 | 67.8 |
| average_vertex_depth | **59.6** | **31.2** | 65.1 | 75.3 |
| average_leaf_depth | **59.6** | **31.2** | 65.1 | 75.3 |
| variance_of_leaves_depths | **79.2** | **66.9** | 85.3 | 89.4 |
| maximum_depth | **77.5** | **58.1** | 82.5 | 92.6 |
| s_shape | **63.8** | **37.8** | 69.5 | 76.5 |
| B_1_index | 7.1 | 14.0 | 6.1 | 4.8 |
| B_2_index | **41.9** | **78.7** | 52.2 | 37.9 |
| maximum_width | -5.2 | -5.2 | 16.4 | 16.4 |
| maxdiff_widths | -8.8 | -8.8 | 17.9 | 17.9 |
| modified_maxdiff_widths | 20.4 | 20.4 | 45.7 | 45.7 |
| max_width_over_max_depth | 16.2 | 18.8 | 13.5 | 13.9 |
| d_index | **42.1** | **15.7** | 49.7 | 61.0 |
| rooted_quartet_index | **36.4** | **53.8** | 42.5 | 23.9 |
| average_ladder | -13.8 | -13.8 | 41.2 | 41.2 |
| ladder_length | -13.8 | -13.8 | 41.2 | 41.2 |
| cherry_index | -1.7 | -1.7 | 12.4 | 12.4 |
| modified_cherry_index | 59.3 | 59.3 | 59.3 | 59.3 |
| IL_number | 59.3 | 59.3 | 59.3 | 59.3 |
| pitchforks | **0.1** | **24.1** | 40.8 | 40.8 |
| four_caterpillars | -2.5 | -2.5 | 16.1 | 16.1 |
| double_cherries | -23.0 | -23.0 | -23.0 | -23.0 |
| total_cophenetic_index | **43.8** | **20.3** | 47.6 | 58.5 |
| diameter | 86.8 | 86.8 | 97.0 | 97.0 |
| area_per_pair_index | 85.8 | 92.8 | 91.3 | 86.2 |
| wiener_index | **28.7** | **42.0** | 42.4 | 31.8 |
| maximum_closeness | 17.7 | 13.7 | 13.2 | 16.9 |
| minimum_farness | 79.4 | 83.7 | 86.8 | 83.1 |
| maximum_farness | 75.4 | 78.2 | 81.7 | 80.8 |
| total_farness | **28.7** | **42.0** | 42.4 | 31.8 |
| minimum_bcent | **0.0** | **53.2** | 53.2 | 0.0 |
| maximum_bcent | 0.1 | 0.1 | 0.1 | 0.1 |
| mean_bcent | 68.7 | 78.5 | 78.7 | 71.2 |
| bcent_variance | **26.0** | **7.4** | 18.0 | 25.2 |
| bcent_root | **0.0** | **70.4** | 70.4 | 53.2 |
| root_imbalance | **46.8** | **20.9** | 29.6 | 46.8 |
| I_root | 20.2 | 16.5 | 29.6 | 46.8 |

The percentile ranks of *T*_1_ and *T*_2_ are highlighted in bold if they differ by more than 10 from each other. This holds for 29 out of 55 indices, including well established and frequently applied indices such as `colless_index`, `sackin_index`, and `total_cophenetic_index`. The root placement hence substantially impacts the tree shape as quantified by these indices. Additionally, we examine the percentile ranks of the unweighted and weighted means. For the unweighted mean, we observe that it can skew the percentile rank towards results for highly unlikely roots. For the `sackin_index`, for example, the percentile rank of the mean is higher than the percentile ranks of *T*_1_, and *T*_2_. This is caused by trees which are rooted at highly unlikely positions but yield higher percentile ranks. The weighted mean can be helpful to balance the percentile ranks of *T*_1_, and *T*_2_, but may be misleading if the percentile ranks of *T*_1_ and *T*_2_ are very distinct. However, calculating the weighted mean is computationally intensive, as the LWRs must be inferred via an exhaustive root search with RootDigger.

Overall, the results illustrate that the impact of the root on certain tree shape indices is so pronounced that it will affect the final results and interpretation of a tree shape study as well as potential downstream analyses. For specific indices, the findings of this case study are only partly in line with our kurtosis-based analysis (see Section 2.1) of all trees in our large-scale evaluation. On the one hand, this illustrates that rooting instability is hard-to-quantify and cannot be captured via a single metric. On the other hand, it also shows, that the actual effects also depend on the specific tree under study.

## 3 Conclusion

In this work, we conduct a large-scale evaluation of 54 tree shape indices on almost 45,000,000 rooted trees obtained from the EvoNAPS and RAxML Grove empirical tree databases. We analyze the results with regard to rooting instability (Section 2.1), correlation with tree size (Section 2.3), and correlation of the indices with each other (Section 2.4).

We find that most tree shape indices exhibit a certain sensitivity to the root position. This behavior is not generally problematic as it may merely reflect the intuition underlying a certain index. In phylogenetics, however, the root position is often uncertain. If the values of an index depend on the root, this inherent uncertainty affects the results of the tree shape analysis.

Our findings further show, that the values of numerous indices do correlate with tree size, sometimes, even so, after applying a normalization technique. In such cases, the index values for different tree sizes can not be compared. By itself, this phenomenon is also not necessarily problematic. The fact that it correlates with trees size, may simply form part of the properties of a particular tree shape index that has a priori not been designed for comparing trees of different sizes. Finally, we observe that numerous indices are strongly correlated with one another. This implies that it suffices to only calculate a subset of the available indices to capture the diverse aspects of a tree’s shape.

Our findings are broadly applicable, as they are based on a huge collection of empirical tree shapes from diverse data sources and organisms. Our results can therefore serve as a guideline for identifying indices with low rooting instability or low correlation to tree size, or to select a subset of indices with low pairwise correlations. However, the indices’ behavior still depends on the specific tree(s) under study, as illustrated by our case study (see Section 2.5).

To circumvent potential pitfalls, we hence recommend conducting the following sanity checks in any tree shape study:

- Evaluate the indices for alternative root positions and assess the impact on the results as already implemented in `treeshapy`
- In case of tree shape indices that are defined on unrooted trees, consider evaluating them directly on the unrooted tree (see Section 2.2)
- Before comparing index values among trees of different sizes, ensure that the correlation between the index and the tree size is low (refer to our results in Section 4.3)
- When evaluating multiple indices, avoid redundancies by checking that the indices’ pairwise correlation is low (refer to our results in Section 2.4). `treeshapy` offers an interface to evaluate the low-correlation index subsets listed in Appendix D.1.

To compute the indices we introduce and release a novel Python library called `treeshapy`. It allows to evaluate all indices in linear time with respect to the number of tips in the tree. As it is based on *ETE3*, it can easily be integrated into existing Python pipelines or frameworks.

We regard tree shape studies as a valuable tool in phylogenetic downstream analysis and tree classification. Through our work, we make tree shape indices more accessible. On the one hand, we provide an easy-to-use Python library with efficient implementations of 56 indices. On the other hand, the above guidelines will facilitate choosing appropriate indices and circumventing interpretation biases.

### 3.1 Outlook and Future Work

Our work provides novel insights but at the same time, raises new questions. Our conclusions are exclusively based on empirical trees. This has the advantage that the results are relevant to practitioners and, at the same time, no bias by a specific tree simulation model is introduced. However, the results may still be biased, as the trees in our databases only cover a tiny fraction of the entire tree space. A large-scale study on broadly sampled tree topologies may provide further insights, but is outside the scope of this work.

Our analyses indicate a substantial rooting instability for many indices (see Section 2.1). Therefore, one may attempt to circumvent rooting altogether and measure tree shapes on unrooted phylogenies instead. Only 12 out of the 56 examined tree shape indices are defined for unrooted trees. Fischer and Liebscher [23] provide a theoretical description of how the shape of unrooted trees can be quantified, but overall, there has been little work on this topic to date.

In Section 2.3 we observe, that none of the known index normalization techniques can completely eliminate the correlation between index values and tree size. This emphasize the need for devising improved normalization approaches. One possibility might be to sample the index distribution for each tree size and normalize the values with the help of these distributions. Since this approach involves sampling a huge amount of trees, it is beyond the scope of this work. Furthermore, eliminating tree size dependency does not constitute the only objective when attempting to devise an appropriate normalization function. Mapping all values of an index to a constant makes that index independent of tree size - but it is clearly not a meaningful normalization technique.

Overall, our results describe the limits of current indices: they are not robust with respect to root placement or they cannot be compared for trees of different sizes. Against this background, we may also ask the question, whether it is possible to design an index without these disadvantages, but that is still able to reflect the tree shape.

Another question is whether multiple indices can be combined to project the tree space into a lower-dimensional space, for example, using PCA. We demonstrate this approach in Appendix D.1, where we perform a PCA in a straightforward manner. The first resulting principal is highly correlated with tree size, the second principal component with `I_root` - which is simply an index exhibiting an exceptionally low correlation with tree size. Our results thus confirm the correlation between most indices and tree size but do not yield a meaningful projection of the tree space into a lower-dimensional space. Investigating whether this is possible with a more sophisticated approach is subject to future work.

## 4 Materials and Methods

In this Section, we first introduce the fundamental notion of a tree shape index. Then, we illustrate the distinct conceptual approaches for characterizing tree shape. Subsequently, we list the indices considered in our study (see Table 4). In addition, we introduce and describe distinct index normalization techniques (see Section 4.1). In Section 4.4, we briefly describe our novel `treeshapy` Python library which we use for all computational experiments. As already mentioned, the main contribution of this study consists in a large-scale as well as comprehensive evaluation that is based on initially unrooted maximum likelihood trees inferred on MSAs from different as well as biologically diverse data sources (described in detail in Section 4.5). In Section 4.6, we describe how we root these un-rooted trees, prior to calculating the corresponding tree shape indices. Additionally, we introduce, and subsequently utilize, the kurtosis (see Section 4.6.1) as a metric to assess the impact of the root position on respective tree shape values. As a last part of this section, we describe the data used in our case study (see Section 4.7).

**Tab. 4.**
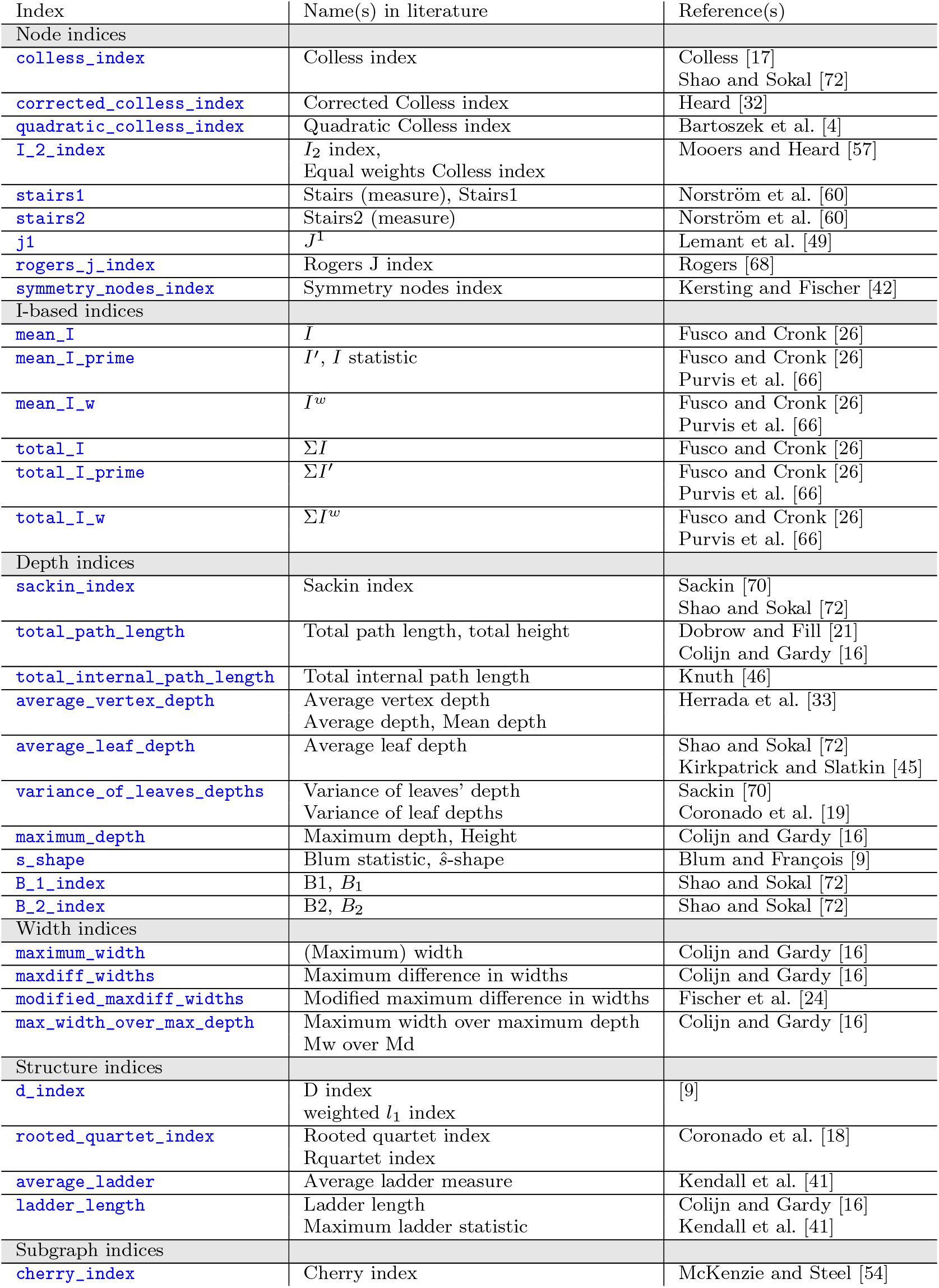
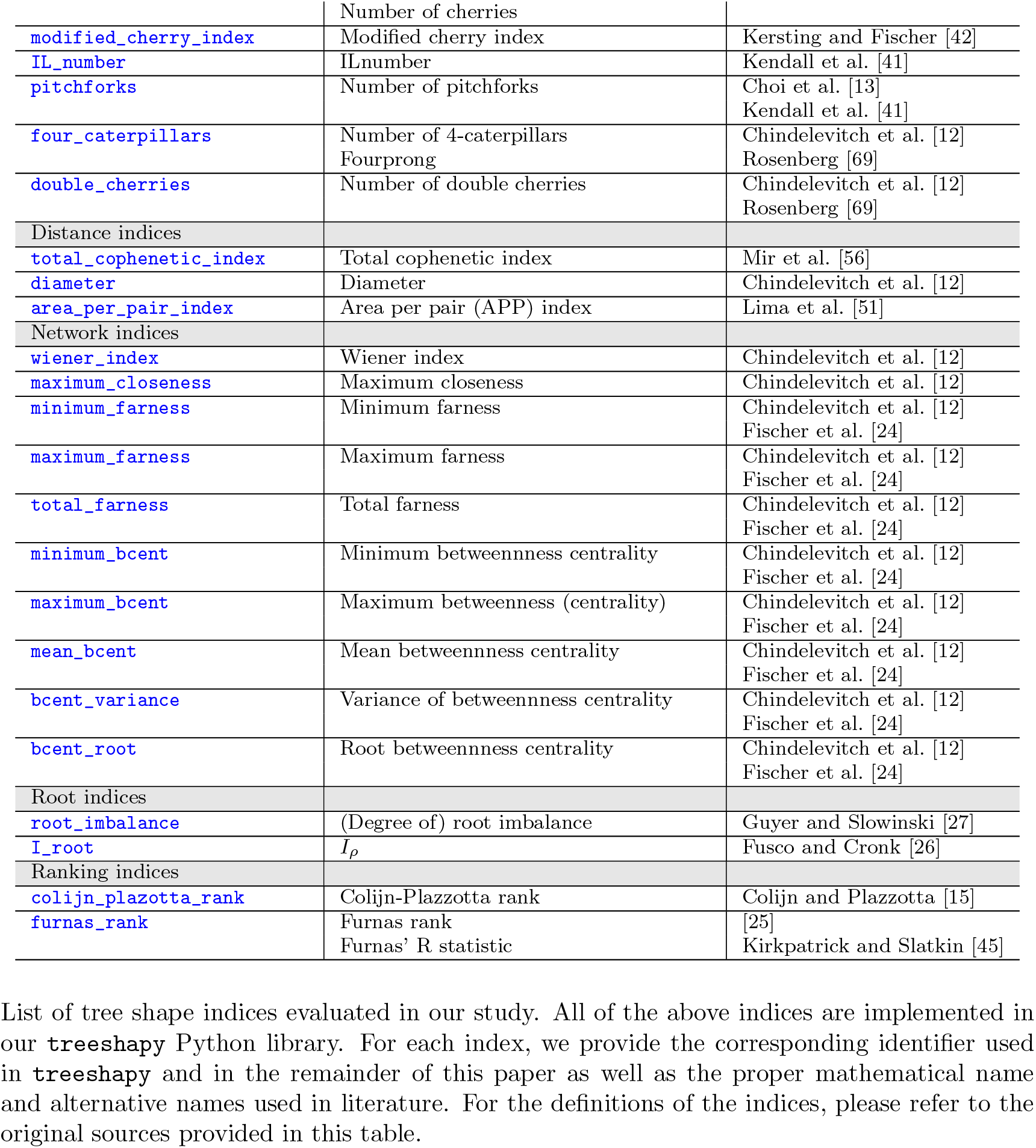
Tree shape indices evaluated in our study.

**Tab. 5.**
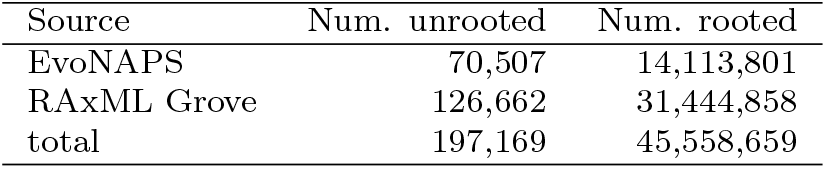
Number of trees under study.

### 4.1 Tree Shape Indices

A tree shape index either assigns an integer or a real valued number to a phylogenetic tree. This facilitates comparing different trees, which would otherwise be difficult due to the high dimensionality of the tree space [38]. The mapping function depends on the specific tree shape characteristics that the respective index intends to capture. Numerous indices focus on tree (im)balance [24].

We classify indices into index categories according to the general approach they adopt for quantifying tree shape. We follow the coarse classification proposed by Janzen and Etienne [38], but further refine it. We illustrate the intuition behind the different approaches and their classification listed below in Fig. 11.

**Fig. 11:**
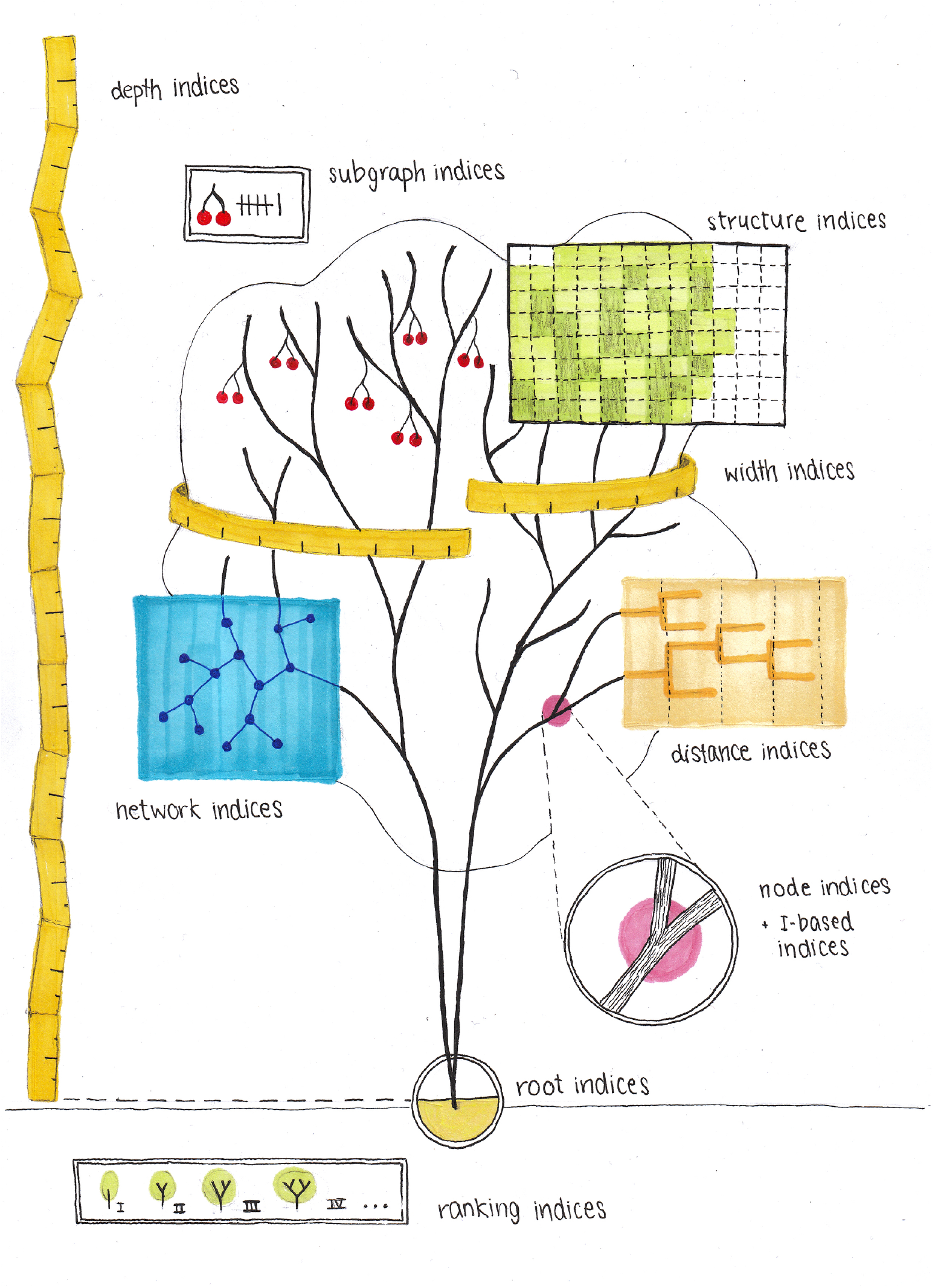
Classes of Tree Shape Indices.

- *Node indices* quantify the balance or shape of a tree via the accumulated properties of its nodes
- *I-based indices* are a subgroup of node indices that are based on the *I*-values [26, 66] of the inner nodes
- *Depth indices* rely on analyzing node depths that correspond to subtree heights
- *Width indices* are based on a tree’s respective widths, that is, the number of nodes exhibiting the same depth
- Under *Structure indices* we group indices that measure the shape of the tree by analyzing the specific sub-structures it contains (for example ladders or induced quartets)
- The values of *subgraph indices* capture the number of occurrences of a certain sub-graph type in the tree (e.g., a cherry)
- *Distance indices* aim to capture tree shape based on the distances of the tree’s leaves to each other
- The idea of *network indices* is to apply concepts from network science for quantifying tree shape
- *Root indices* exclusively focus on the balance of the root node
- *Ranking indices* are based on ordering all topologies of the same size according to their shape. The resulting value corresponds to the rank of the tree in this order

The selection of indices in our study (see Table 4) is based on the surveys by Fischer et al. [24] and Janzen and Etienne [38]. However, we ensure that our results are manageable even when evaluating the indices on a large amount of trees - which is the main focus of our work. To this end, we limit ourselves to indices that capture the properties of tree topologies and disregard those that also take branch lengths into account (e.g., treeness [2]). Branch length estimates further go along with additional uncertainties [50, 71] which carry on to any tree shape analysis involving them. Furthermore, we omit indices that require branching times (e.g., crown age [38]) or additional parameters (e.g., clades of size *x* [12]).

Note that we exclude rank indices from most of our analyses. This is because they yield very large integer values that are computationally difficult to analyze. Doing so would require the use of a library for multi-precision arithmetic, which is beyond the scope of this work. Furthermore, we regard the rank indices as being of secondary importance, since the large integer values they return are also difficult to handle in practice. Nevertheless, the rank indices *can* be computed using our Python library for trees of any size (see Section 4.4).

Please note that the assignment of indices to the respective classes is not necessarily unique. It merely serves to better structure our analyses and maintain an overview of the numerous indices. `bcent_root`, for example, is based on betweenness centrality which is a concept from network science. For this reason, we consider `bcent_root` to be a network index. At the same time, it could also have been classified as a root index, since it corresponds to the betweenness centrality of the root.

List of tree shape indices evaluated in our study. All of the above indices are implemented in our `treeshapy` Python library. For each index, we provide the corresponding identifier used in `treeshapy` and in the remainder of this paper as well as the proper mathematical name and alternative names used in literature. For the definitions of the indices, please refer to the original sources provided in this table.

### 4.2 Sampling Tree Topologies

For a fixed value *n*, the corresponding tree space comprises all possible tree topologies with *n* leaves. When analyzing a tree shape index, the distribution of index values within this tree space constitutes a relevant criterion. However, the size of the tree space grows super-exponentially as a function of *n*. Therefore, even for relatively small values of *n*, it is infeasible to determine the corresponding distribution. Instead, one approximates it by sampling topologies from the tree space using a probabilistic model. Various models have been introduced to this end [1, 30, 28, 24], but we restrict ourselves to the *Yule model* [76] in the context of this work. This model is simple in the sense that it yields no parameters except the tree size. Further, it has frequently been used in previous works.

### 4.3 Normalizing Tree Shape Indices

In the following, we introduce three different normalization techniques, which can be applied to absolute tree shape index values. All normalization approaches aim to eliminate (or at least reduce) the dependency of the index values on the tree size in terms of number of leaves to yield the corresponding values more comparable (see Section 2.3).

Let *I* be a tree shape index and let *T* be a rooted tree with *n* leaves.

#### 4.3.1 Normalization by Number of Tips

By *I*_rt_(*T* ) we denote the value that *I* assigns to *T* normalized by the number of tips *n*. We define

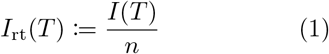

This normalization technique can be applied to all indices.

#### 4.3.2 Normalization by Maximum Value

Further, let *I*_max_(*n*) (*I*_min_(*n*)) be the maximum (minimum) value of *I* over all possible trees with *n* leaves. Note that proven closed forms of these values are only available for 19 out of the 56 indices under study (see Table 6). For the remaining indices, this normalization approach is not possible. By *I*_rm_(*T* ), we denote the normalized value that *I* assigns to *T* . Following Fischer et al. [24], we define

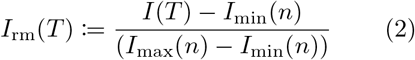

It holds that *I*_rm_(*T* ) ∈ [0, 1].

In binary trees of size *n* but for which *n* is *not* a power of 2, the maximum value may never be attained [32]. This potentially biases the results of this normalization technique [24].

#### 4.3.3 Normalization by Expected Value under the Yule Model

Let E^*I*^ (*n*) be the expected value of *I* for a tree with *n* leaves that was generated under the Yule model. Note that the following normalization approach is only possible for 11 out of 55 for indices, for which E^*I*^ (*n*) is known (see Table 6). By *I*_ry_(*T* ), we denote the value that *I* assigns to *T* normalized by this expected value. Following Blum and François [9] and Janzen and Etienne [38], we define

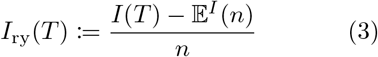

Kirkpatrick and Slatkin [45] mention a similar normalization 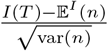 which we do not investigate in greater detail. This is because, their normalization also requires var(*n*), that is, the variance under the Yule model, such that it is defined for even fewer indices.

### 4.4 The treeshapy Python library

We conduct our computational experiments using our novel `treeshapy` Python library which implements all 56 tree shape indices listed in Table 4. The evaluation of each index requires linear time with respect to the number of tips *n* in the input tree. In the appendix, we present a detailed performance benchmark (see Appendix A.1). Additionally, we describe, how we verify the correctness of `treeshapy` (see Appendix A.2).

`treeshapy` is based on the tree data structure used in *ETE3* [36]. This tree library is widely used in computational phylogenetics and therefore facilitates integrating `treeshapy` into larger phylogenetic data analysis pipelines. We use the tree node attributes as provided by ETE3 to store intermediate results.

As outlined in the preceding Section 4.3, the absolute value of a tree shape index can (potentially) be normalized via the maximum value, the expected value under the Yule model, or the number of tips (see Section 4.3). `treeshapy` generally supports all normalization options. However, the minimum value, the maximum value, and the expected value under the Yule model are unknown for numerous indices. Hence, the corresponding normalization is not feasible. In Table 6 of the appendix we list which normalization technique is implemented for which index.

**Tab. 6.** Additional information about the indices.

| Index | 1 | 2 | 3 | 4 | 5 | 6 | 7 | 8 |
| --- | --- | --- | --- | --- | --- | --- | --- | --- |
| Node indices |  |  |  |  |  |  |  |  |
| colless_index | ✓ | ✓ | I | ✗ | ✓ | ✗ | ✓ | ✗ |
| corrected_colless_index | ✓ | ✓ | I | ✗ | ✓ | ✗ | ✓ | ✗ |
| quadratic_colless_index | ✓ | ✓ | I | ✗ | ✓ | ✗ | ✓ | ✗ |
| I_2_index | ✓ | ✓ | I | ✗ | ✗ | ✗ | ✗ | ✗ |
| stairs1 | ✓ | ✓ | I | ✗ | ✓ | ✗ | ✗ | ✗ |
| stairs2 | ✓ | ✓ | B | ✗ | ✗ | ✗ | ✗ | ✗ |
| j1 | ✗ | ✓ | - | ✓ | ✗ | ✗ | ✗ | ✗ |
| rogers_j_index | ✓ | ✓ | I | ✗ | ✓ | ✗ | ✗ | ✗ |
| symmetry_nodes_index | ✓ | ✓ | I | ✗ | ✓ | ✗ | ✗ | ✗ |
| I-based indices |  |  |  |  |  |  |  |  |
| mean_I | ✓ | ✗ | I | ✗ | ✗ | ✗ | ✗ | ✗ |
| mean_I_prime | ✓ | ✓ | I | ✗ | ✗ | ✗ | ✗ | ✗ |
| mean_I_w | ✓ | ✗ | I | ✗ | ✗ | ✗ | ✗ | ✗ |
| total_I | ✓ | ✗ | I | ✗ | ✗ | ✗ | ✗ | ✗ |
| total_I_prime | ✓ | ✗ | I | ✗ | ✗ | ✗ | ✗ | ✗ |
| total_I_w | ✓ | ✗ | I | ✗ | ✗ | ✗ | ✗ | ✗ |
| Depth indices |  |  |  |  |  |  |  |  |
| sackin_index | ✓ | ✓ | I | ✓ | ✓ | ✓ | ✓ | ✗ |
| total_path_length | ✓ | ✓ | I | ✓ | ✓ | ✓ | ✗ | ✗ |
| total_internal_path_length | ✓ | ✓ | I | ✓ | ✓ | ✓ | ✗ | ✗ |
| average_vertex_depth | ✓ | ✓ | I | ✓ | ✓ | ✓ | ✗ | ✗ |
| average_leaf_depth | ✓ | ✓ | I | ✓ | ✓ | ✓ | ✓ | ✗ |
| variance_of_leaves_depths | ✓ | ✓ | I | ✓ | ✗ | ✓ | ✓ | ✗ |
| maximum_depth | ✓ | ✓ | I | ✓ | ✓ | ✓ | ✗ | ✗ |
| s_shape | ✓ | ✓ | I | ✓ | ✗ | ✓ | ✗ | ✗ |
| B_1_index | ✓ | ✓ | B | ✓ | ✗ | ✗ | ✗ | ✗ |
| B_2_index | ✓ | ✓ | B | ✓ | ✓ | ✓ | ✓ | ✗ |
| Width indices |  |  |  |  |  |  |  |  |
| maximum_width | ✓ | ✓ | B | ✓ | ✗ | ✓ | ✗ | ✗ |
| maxdiff_widths | ✓ | ✗ | - | ✓ | ✗ | ✓ | ✗ | ✗ |
| modified_maxdiff_widths | ✓ | ✗ | B | ✓ | ✗ | ✓ | ✗ | ✗ |
| max_width_over_max_depth | ✓ | ✓ | B | ✓ | ✗ | ✗ | ✗ | ✗ |
| Structure indices |  |  |  |  |  |  |  |  |
| d_index | ✓ | ✗ | - | ✓ | ✗ | ✗ | ✗ | ✗ |
| rooted_quartet_index | ✓ | ✓ | B | ✓ | ✗ | ✓ | ✓ | ✗ |
| average_ladder | ✗ | ✓ | - | ✗ | ✗ | ✗ | ✗ | ✗ |
| ladder_length | (✓) | ✓ | - | ✗ | ✗ | ✗ | ✗ | ✗ |
| Subgraph indices |  |  |  |  |  |  |  |  |
| cherry_index | ✓ | ✓ | - | ✓ | ✓ | ✓ | ✓ | ✗ |
| modified_cherry_index | ✓ | ✗ | - | ✗ | ✓ | ✗ | ✗ | ✗ |
| IL_number | (✓) | ✓ | - | ✓ | X | X | X | X |
| pitchforks | (✓) | ✓ | - | ✓ | X | X | X | X |
| four_caterpillars | (✓) | ✓ | - | ✓ | X | X | X | X |
| double_cherries | (✓) | ✓ | - | ✓ | X | X | X | X |
| Distance indices |  |  |  |  |  |  |  |  |
| total_cophenetic_index | ✓ | ✓ | I | ✓ | ✓ | ✓ | ✓ | X |
| diameter | (✓) | ✓ | - | ✓ | X | ✓ | X | ✓ |
| area_per_pair_index | ✓ | ✓ | - | ✓ | X | X | ✓ | ✓ |
| Network indices |  |  |  |  |  |  |  |  |
| wiener_index | (✓) | ✓ | - | ✓ | X | X | X | ✓ |
| maximum_closeness | X | ✓ | - | ✓ | X | X | X | ✓ |
| minimum_farness | (✓) | X | - | ✓ | X | X | X | ✓ |
| maximum_farness | (✓) | X | - | ✓ | X | X | X | ✓ |
| total_farness | (✓) | X | - | ✓ | X | X | X | ✓ |
| minimum_bcent | (✓) | X | - | ✓ | X | X | X | ✓ |
| maximum_bcent | (✓) | ✓ | - | ✓ | X | X | X | ✓ |
| mean_bcent | (✓) | X | - | ✓ | X | X | X | ✓ |
| bcent_variance | (✓) | X | - | ✓ | X | X | X | ✓ |
| bcent_root | (✓) | X | - | ✓ | X | X | X | X |
| Root indices |  |  |  |  |  |  |  |  |
| root_imbalance | (✓) | ✓ | - | X | ✓ | X | X | X |
| I_root | (✓) | X | - | X | ✓ | X | X | X |
| Ranking indices |  |  |  |  |  |  |  |  |
| colijn_plazotta_rank | ✓ | X | I | X | X | X | X | X |
| furnas_rank | ✓ | X | B | X | ✓ | X | X | X |

1 - implemented by Fischer et al. [24]
2 - implemented by Janzen and Etienne [38]
3 - index type (B balance, I imbalance, - none) according to Fischer et al. [24]
4 - defined on arbitrary trees
5 - can be normalized by maximum for binary trees
6 - can be normalized by maximum for arbitrary trees
7 - can be normalized by expected value under the Yule model (binary trees only)
8 - defined for unrooted trees

Some tree shape indices are only defined on strictly binary (bifurcating) trees, while others can be evaluated on arbitrary (multifurcating) trees (see Table 6 in the appendix). The minima and maxima may also differ between bifurcating and multi-furcating trees. This affects the values when they are normalized using the maximum. Therefore, `treeshapy` offers two modes, one for binary and one for arbitrary trees. This ensures that the normalization works properly and that indices are only evaluated on the tree types for which they have been defined. Note that the trees we use in our experiments are all strictly binary. Hence, we perform all evaluations presented here in binary tree mode. We have nevertheless, fully implemented and extensively tested the arbitrary tree mode as well (see Appendix A.2).

### 4.5 Tree Data

We use `treeshapy` to perform a large-scale evaluation of tree shape indices on a total of almost 200,000 unrooted trees obtained from the *EvoNAPS* and *RAxML Grove* databases.

EvoNAPS is a comprehensive phylogenetic database containing empirical data. To build the database, [67] collected multiple sequence alignments (MSAs) from three different sources: the online repository BenchmarkAlignments [48], the OrthoMaM database [22], the PANDIT database [75], and TreeBase [62]. For each MSA, a maximum likelihood tree was inferred using IQ Tree [55]. From EvoNAPS we use a total of more than 70,000 phylogenetic trees.

RAxML Grove [37] comprises more than 120,000 trees inferred on datasets which were analyzed on the RAxML or RAxML-NG webservers. The trees under study vary in size (see Fig. 12 for the corresponding tree size distribution). Note that we omit trees with more than 2,500 taxa since they yield a large number of rooted trees combined with increased runtimes for index evaluation.

**Fig. 12:**
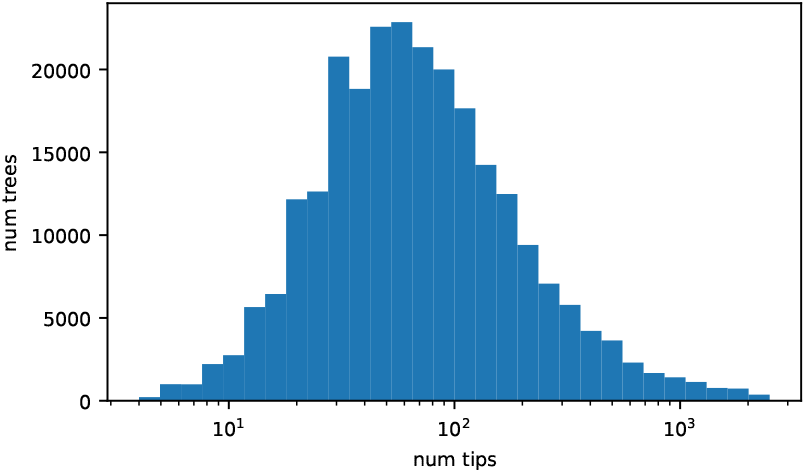
Distribution of empirical tree sizes in our study. Note the logarithmic scale along the x-axis.

### 4.6 Rooting Trees

Most tree shape indices we implemented and examine require a rooted input tree [24]. However, trees obtained via phylogenetic inference methods, including the EvoNAPS and RAxML Grove trees, are typically unrooted [55, 47]. These unrooted trees can be rooted via different downstream approaches, such as outgroup rooting [74] or midpoint rooting [73]. The *RootDigger* tool uses a non-reversible Markov model of character substitution to determine the most likely position as well as the corresponding placement probability of the root. IQ-TREE 2 [55] also allows to compute such root placement probabilities, called *rootstrap supports* [59]. What all these down-stream rooting approaches have in common though, is an inherent uncertainty regarding the root placement [6, 44, 34].

This inherent uncertainty naturally leads to the question to which extent the respective index values depend on the root position. In other words: How does the potentially erroneous or uncertain root position affect the results of a tree shape analysis? In the following we refer to this concept as *rooting instability*.In the following experiments, we root the unrooted EvoNAPS and RAxML Grove trees on every possible branch. For an unrooted tree *T* with *n* leaves, rooting on every possible branch yields a set ℛ (*T* ) of 2*n* − 3 rooted trees. For all rooted trees we obtain in this way, we calculate all implemented tree shape indices. We use the kurtosis Kurt as a metric to capture rooting instability.

Note that, `diameter`, `area_per_pair` as well as *all* network indices (except `bcent_root`) can be calculated on unrooted trees. However, this does not mean that these indices will yield the same index value on all rooted trees that have been constructed (as outlined above) from the unrooted reference tree. This is because, rooting a tree involves inserting an additional node — the root — into a branch of the unrooted tree. This modification can affect the index results, especially if the index can also be calculated on an unrooted tree. In Section 2.2, we examine the behavior of these indices in greater detail.

#### 4.6.1 Kurtosis

Let *X* be a random variable with mean *µ* and standard deviation *σ*. The Kurtosis Kurt(*X*) is defined as the standardized fourth moment:

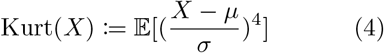

[61] In order to measure the rooting instability of an unrooted tree *T* and an index *I*, we set Kurt(*I*(*T* )) := Kurt(*I*(*R*)), where *R* is a random variable that follows a uniform distribution on ℛ (*T* ) (see Fig. 13).

**Fig. 13:**
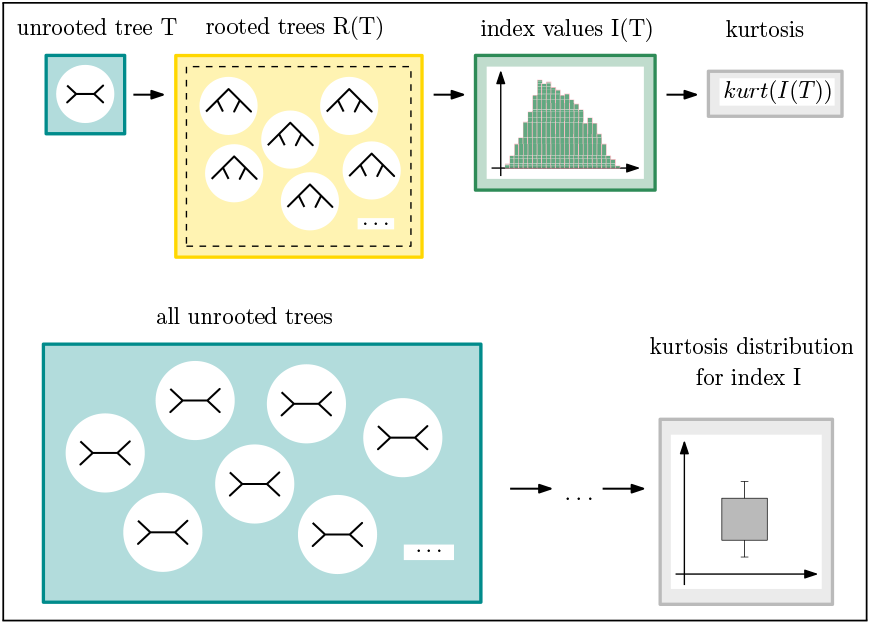
Experimental setup to determine the kurtosis distributions.

**Fig. 14:**
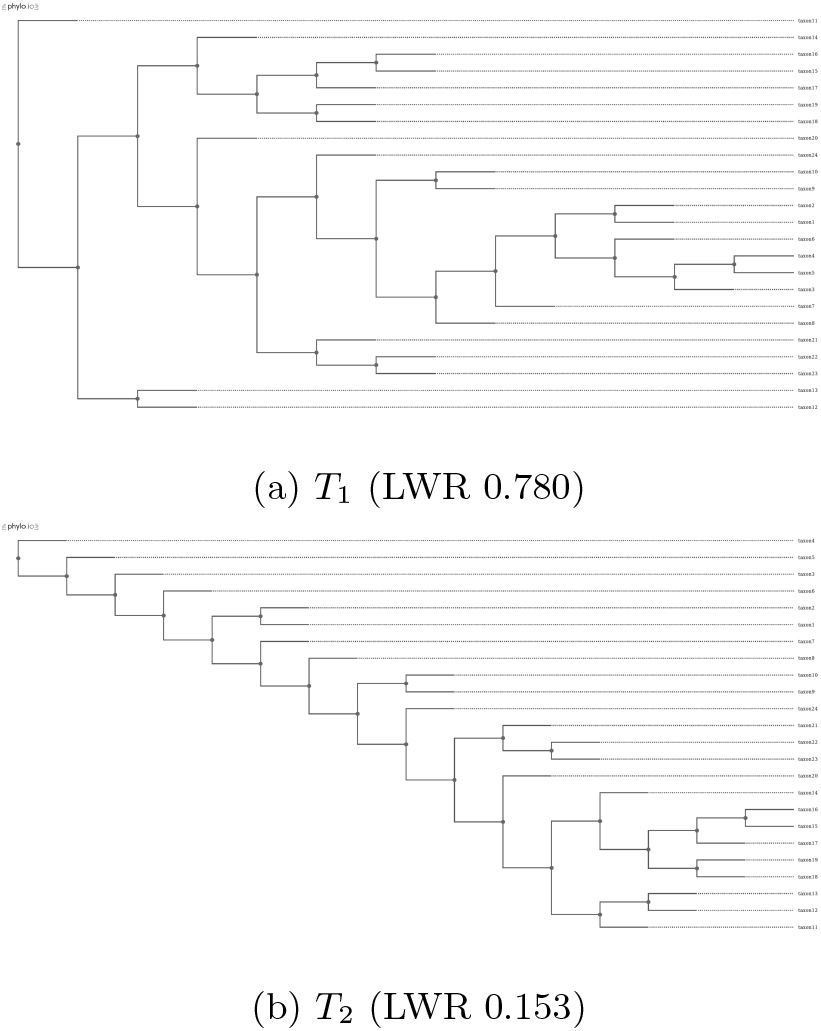
Receptor tree used in the case study. *T*_1_ is rooted in the branch with the highest LWR, *T*_2_ in the branch with the second highest LWR.

The kurtosis of a uniform (normal) distribution is 1.8 (3). If we observe Kurt(*I*(*T* )) *>* 3, the respective index value distribution has heavier tails and higher peaks than the normal distribution, which indicates a higher rooting instability. If we observe Kurt(*I*(*T* )) *<* 3 instead, the index value distribution is flatter and has less pronounced peaks which we interpret as indicating lower rooting instability [20]. Interpreting the kurtosis in this manner is based on the assumption that rooting stability is predominantly affected by the extreme values of the index.

Further note that, due to its standardization, the kurtosis of a distribution is independent of the nominal values. We can hence directly compare the kurtosis of different trees and indices, despite the fact that the index values depend on the tree size and the distinct indices may differ in their value ranges.

### 4.7 Case Study

For our case study, we select an MSA of length 73 with 24 DNA sequences from a specific receptor gene in humans, dromedary, sheep and dolphins [52] (Additional File 4, TreeBase ID S19396). We infer a Maximum Likelihood tree on this MSA which we call receptor tree in the following. The exhaustive root placement search mode in RootDigger assigns a *likelihood weight ratio (LWR)* to each branch of the tree. This per-branch LWR quantifies the probability of placing the root onto the specific branch Bettisworth and Stamatakis [6].

For the receptor tree, we obtain a maximum LWR of 0.780 for placing the root on a specific branch from the exhaustive root search with RootDigger. However, for a distinct, alternative branch, we obtain a root LWR of 0.153. For this dataset there is hence no clear root placement signal. Instead, there are two branches for which the root placement probability is not negligible. Further, these two alternative root placement branches are far away from each other. They are 12 nodes apart in the unrooted tree, which yields a diameter of 16. Hence, the two resulting rooted tree topologies differ substantially.

For our experiments, we root the un-rooted receptor tree at every possible branch. Thereby, we obtain a set of rooted trees on which we calculate the tree shape indices under study. We denote the trees with roots located on the two branches with high LWRs by *T*_1_ (LWR = 0.780), and *T*_2_ (LWR = 0.153).

### 4.8 Data Availability

`treeshapy` is available on Github and can be installed via pip. Our experiments are also available on Github. Additionally, all data can be obtained from Zenodo.

## Acknowledgement

We would like to thank Franzsika Reden for the initial experiments on the EvoNAPS database, Tanja Stadler for a helpful exchange and Alexander Jordan for sharing his expertise to find a metric for measuring rooting instability.

Luise Häuser and Alexandros Stamatakis are financially supported by the Klaus Tschira Foundation, and by the European Union (EU) under Grant Agreement No 101087081 (CompBiodiv-GR).

---

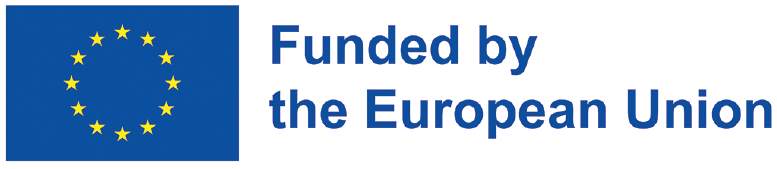

## A treeshapy

### A.1 Performance Benchmark

In the following, we analyze the performance of `treeshapy` by benchmarking the runtimes of the index computations with our large-scale evaluation on EvoNAPS and RAxML Grove trees. We perform all evaluations are on an AMD EPYC 7413 24-Core Processor (1.5 GHz) (512 GB RAM, Ubuntu 24.04). Note that we have not yet parallelized `treeshapy`.

For each rooted tree, we measure the time required to evaluate all implemented indices without normalization. Many of the indices considered require node properties such as `depth` or `clade_size` (see Table 7). In the benchmarking version of `treeshapy`, we therefore pre-compute these node properties prior to computing the actual indices. The node properties can then be accessed in constant time whenever required during index calculation. The respective pre-computation time is included in the overall runtimes we provide. For all trees of the same size, we calculate the mean runtime. The results are shown in Fig. 15.

**Tab. 7.** Dependencies of the indices.

| Index | Dependencies |
| --- | --- |
| Node indices |  |
| `colless_index` | `clade_size` |
| `corrected_colless_index` | `clade_size`, `colless_index` |
| `quadratic_colless_index` | `clade_size` |
| `I_2_index` | `clade_size` |
| `stairs1` | `clade_size`, `rogers_j_index` |
| `stairs2` | `clade_size` |
| `j1` | `clade_size` |
| `rogers_j_index` | `clade_size` |
| `symmetry_nodes_index` |  |
| I-based indices |  |
| `mean_I` | `clade_size` |
| `mean_I_prime` | `clade_size` |
| `mean_I_w` | `clade_size` |
| `total_I` | `clade_size` |
| `total_I_prime` | `clade_size` |
| `total_I_w` | `clade_size` |
| Depth indices |  |
| `sackin_index` | `depth` |
| `total_path_length` | `depth` |
| `total_internal_path_length` | `depth` |
| `average_vertex_depth` | `depth` |
| `average_leaf_depth` | `depth` |
| `variance_of_leaves_depths` | `depth` |
| `maximum_depth` | `depth` |
| `s_shape` | `clade_size` |
| `B_1_index` |  |
| `B_2_index` |  |
| Width indices |  |
| `maximum_width` | `depth` |
| `maxdiff_widths` | `depth` |
| `modified_maxdiff_widths` | `depth` |
| `max_width_over_max_depth` | `maximum_width`, `maximum_depth`, `depth` |
| Structure indices |  |
| `d_index` | `clade_size` |
| `rooted_quartet_index` | `clade_size` |
| `average_ladder` | `ladder_lengths` |
| `ladder_length` | `ladder_lengths` |
| Subgraph indices |  |
| `cherry_index` |  |
| `modified_cherry_index` | `cherry_index`, `clade_size` |
| `IL_number` |  |
| `pitchforks` | `clade_size` |
| `four_caterpillars` | `clade_size` |
| `double_cherries` | `clade_size` |
| Distance indices |  |
| `total_cophenetic_index` | `clade_size` |
| `diameter` | `depth` |
| `area_per_pair_index` | `depth`, `clade_size`, `sackin_index`, `total_cophenetic_index` |
| Network indices |  |
| `wiener_index` | `nodes_below` |
| `maximum_closeness` | `nodes_below`, `farness` |
| `minimum_farness` | `nodes_below`, `farness` |
| `maximum_farness` | `nodes_below`, `farness` |
| `total_farness` | `nodes_below`, `farness` |
| `minimum_bcent` | `nodes_below`, `bcent` |
| `maximum_bcent` | nodes_below, bcent |
| `mean_bcent` | nodes_below, bcent |
| `bcent_variance` | nodes_below, bcent |
| `bcent_root` | nodes_below, bcent |
| Root indices |  |
| `root_imbalance` | clade_size |
| `I_root` | clade_size |
| Ranking indices |  |
| `colijn_plazotta_rank` |  |
| `furnas_rank` | clade_size |

Dependencies on node properties (grey) and other indices (blue) which we exploit in **treeshapy** to speed up computations

**Tab. 8.** Studies applying tree shape indices.

| Quote | Empirical Trees | Objective | Indices |
| --- | --- | --- | --- |
| [8] | 37 trees inferred on mitochondrial data of different human populations | detect matrilineal fertility inheritance | `mean_I_prime` |
| [10] (Section 5) | 1 tree inferred on HIV-1 data | measure compatibility with the Yule model | `sackin_index`, `D_index`, (number of subtrees of fixed size) |
| [65] | 62 RNA virus trees | match modes of transmission and pathogenesis with tree shape | `colless_index`, `sackin_index`, `average_path_length`, `B_1_index`, `B_2_index`, `total_I`, `mean_I` (mean path length, variance of path lengths, $I_10$, newly introduced kernel trick) |
| [16] | Simulated Data and Trees inferred with BEAST on 2 datasets from outbreaks of tuberculosis | Determine transmission patterns in disease outbreaks | `corrected_colless_index`, `sackin_index`, `il_number`, `ladder_length`, `maximum_width`, `max_width_over_max_depth`, `maxdiff_widths`, `cherry_index`, `stairs1`, `stairs2` |
| [51] | 8,999 trees from Tree-Base | Introduction of the `area_per_pair_index`, comparison to other indices | `area_per_pair_index`, `sackin_index`, `total_cophenetic_index` |
| [5] | HIV datasets | Determine evolutionary regime / selection type in viral trees | `cherry_index`, `pitchforks`, `colless_index`, `sackin_index`, `maximum_height`, `maximum_width`, betweenness centrality, closeness centrality, eigenvector centrality, `diameter`, four metrics based on laplacian spectral properties |
| [40] | 2 previously published trees for HIV data | Train a classifier distinguishing between pathogen phylogenetic trees with structured and unstructured host populations | `cherry_index`, `colless_index`, `sackin_index`, `total_cophenetic_index`, `ladder_length`, `maximum_width`, `maximum_depth`, `max_width_over_max_depth` |
| [11] | 108 animal trees, 110 plant trees | Relationship of tree shape and structure of plant-pollinator interaction networks | `colless_index` |
| [29] | 50 paleontological trees, 50 neontological trees (previously published) | Testing whether paleontological trees are less balanced than neontological trees | `colless_index` |
| [7] | 378 trees from TreeBASE, 14,509 trees from OrthoMaM, 77,843 trees from PhylomeDB, 85,581 trees from HOGENOM | Compare various balance indices in terms of how good they perform in distinguishing trees by their origins | colless_index, sackin_index, cherry_index, variance_of_leaves_depths, B_1_index, B_2_index |
| [31] | Subtrees of a primate phylogeny | Examine diversity skewness | mean_I |
| [63] | Tree inferred with BEAST on tuberculosis data | Detect effects of non-exponential infectious periods on pathogens' transmission dynamics | colless_index, sackin_index, cherry_index, pitchforks, ladder_length, stairs2, il_number |
| [53] | 10 species trees published in previous studies | Relationship of radiation (after isolation of a species) and tree balance | (normalized) sackin_index |
| [35] | Language Tree from ethnologue | Draw conclusions about rates of origination and extinction by studying tree balance | mean_I |
| [60] | 19 HIV datasets | Test PhyloTempo library | stairs1, staris2, sackin_index, cherry_index, beta[1] |
| [64] | 6 RNA virus trees | Connection of tree balance to the extend of virus-host interaction | average_vertex_depth, mean_I_prime, mean topological distance |
| [12] | Trees inferred on data from 3 different viruses with RAxML-NG, additional previously published HIV and Influenza trees | Examine how the tree shape indices distinguish between different viruses and epidemiological scenarios | cherry_index, pitchforks, double_cherries, four_caterpillars, colless_index, sackin_index, maximum_depth, maximum_width, stairs1, maxdiff_widths, diameter, wiener_indexclades of size x, mean pairwise distance, spectral properties, distance Laplacian spectral properties, node properties from network science |

**Tab. 9.** Pearson correlation of index values with tree size.

| index | abs. | rel.(tips) | rel.(max) | rel.(yule) |
| --- | --- | --- | --- | --- |
| colless_index | 0.85 | 0.54 | -0.34 | 0.49 |
| corrected_colless_index | -0.34 | -0.14 | -0.34 | -0.13 |
| quadratic_colless_index | 0.69 | 0.76 | -0.29 | 0.74 |
| I_2_index | -0.09 | -0.21 |  |  |
| stairs1 | -0.03 | -0.25 | -0.03 |  |
| stairs2 | 0.03 | -0.23 |  |  |
| j1 | -0.28 | -0.2 |  |  |
| rogers_j_index | 1.0 | 0.02 | -0.03 |  |
| symmetry_nodes_index | 1.0 | 0.03 | -0.02 |  |
| mean_I | -0.07 | -0.23 |  |  |
| mean_I_prime | -0.06 | -0.24 |  |  |
| mean_I_w | -0.07 | -0.22 |  |  |
| total_I | 0.98 | -0.03 |  |  |
| total_I_prime | 0.98 | -0.03 |  |  |
| total_I_w | 0.97 | -0.04 |  |  |
| sackin_index | 0.89 | 0.6 | -0.3 | 0.49 |
| total_path_length | 0.89 | 0.6 | -0.3 |  |
| total_internal_path_length | 0.88 | 0.6 | -0.3 |  |
| average_vertex_depth | 0.6 | -0.41 | -0.3 |  |
| average_leaf_depth | 0.6 | -0.41 | -0.3 | -0.21 |
| variance_of_leaves_depths | 0.33 | -0.05 |  | -0.02 |
| maximum_depth | 0.64 | -0.39 | -0.31 |  |
| s_shape | 0.99 | 0.14 |  |  |
| B_1_index | 0.99 | 0.07 |  |  |
| B_2_index | 0.03 | -0.28 | -0.09 | 0.32 |
| maximum_width | 0.89 | -0.35 |  |  |
| maxdiff_widths | 0.64 | -0.33 |  |  |
| modified_maxdiff_widths | 0.7 | -0.33 |  |  |
| max_width_over_max_depth | 0.4 | -0.22 |  |  |
| d_index | 0.59 | -0.43 |  |  |
| rooted_quartet_index | 0.68 | 0.76 |  | -0.71 |
| average_ladder | 0.0 | -0.23 |  |  |
| ladder_length | 0.39 | -0.25 |  |  |
| cherry_index | 0.99 | 0.07 | 0.09 | 0.07 |
| modified_cherry_index | 0.98 | -0.07 | -0.09 |  |
| IL_number | 0.98 | -0.07 |  |  |
| pitchforks | 0.99 | 0.02 |  |  |
| four_caterpillars | 0.97 | -0.05 |  |  |
| double_cherries | 0.87 | -0.01 |  |  |
| total_cophenetic_index | 0.71 | 0.78 | -0.29 | 0.74 |
| diameter | 0.69 | -0.41 |  |  |
| area_per_pair_index | 0.73 | -0.41 |  | -0.01 |
| wiener_index | 0.85 | 0.94 |  |  |
| maximum_closeness | -0.13 | -0.07 |  |  |
| minimum_farness | 0.93 | 0.66 |  |  |
| maximum_farness | 0.81 | 0.72 |  |  |
| total_farness | 0.85 | 0.94 |  |  |
| minimum_bcent | 0.9 | 0.02 |  |  |
| maximum_bcent | 0.91 | 1.0 |  |  |
| mean_bcent | 0.93 | 0.73 |  |  |
| bcent_variance | 0.78 | 0.85 |  |  |
| bcent_root | 0.24 | 0.1 |  |  |
| root_imbalance | 0.12 | -0.27 | 0.08 |  |
| I_root | 0.08 | -0.24 | 0.08 |  |

The cells are colored according to their values ( $< 0.1$ : green, $[0.1, 0.3]$ : yellow, $[0.3, 0.6]$ : orange, $> 0.6$ : red). If a cell is gray, the normalization technique is not applicable for the respective index (see Section 4.3).

**Fig. 15:**
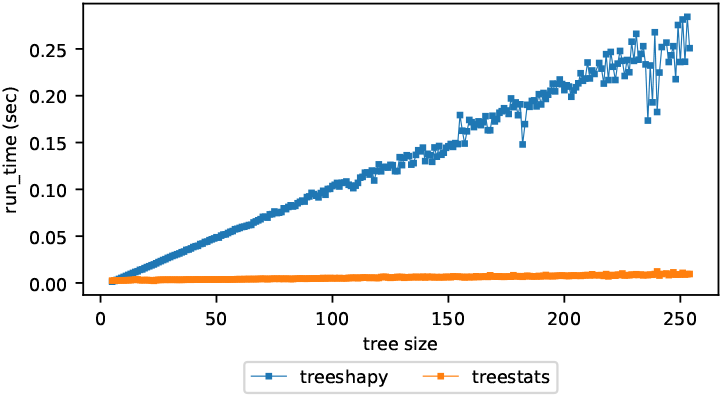
Runtimes of treeshapy **and** treestats compared. The x-axis corresponds to the tree size in terms of number of tips. The y-axis indicates the runtime in seconds. For each tree, we measure the time required to compute all implemented tree shape indices (including the afore-mentioned pre-computation time in the case of `treeshapy`). We bin the trees based on their size in bins of range 10 and provide the mean runtime. The plot illustrates, that index calculations require linear time both using `treeshapy` and `treestats`. However the latter is substantially faster.

**Fig. 16:**
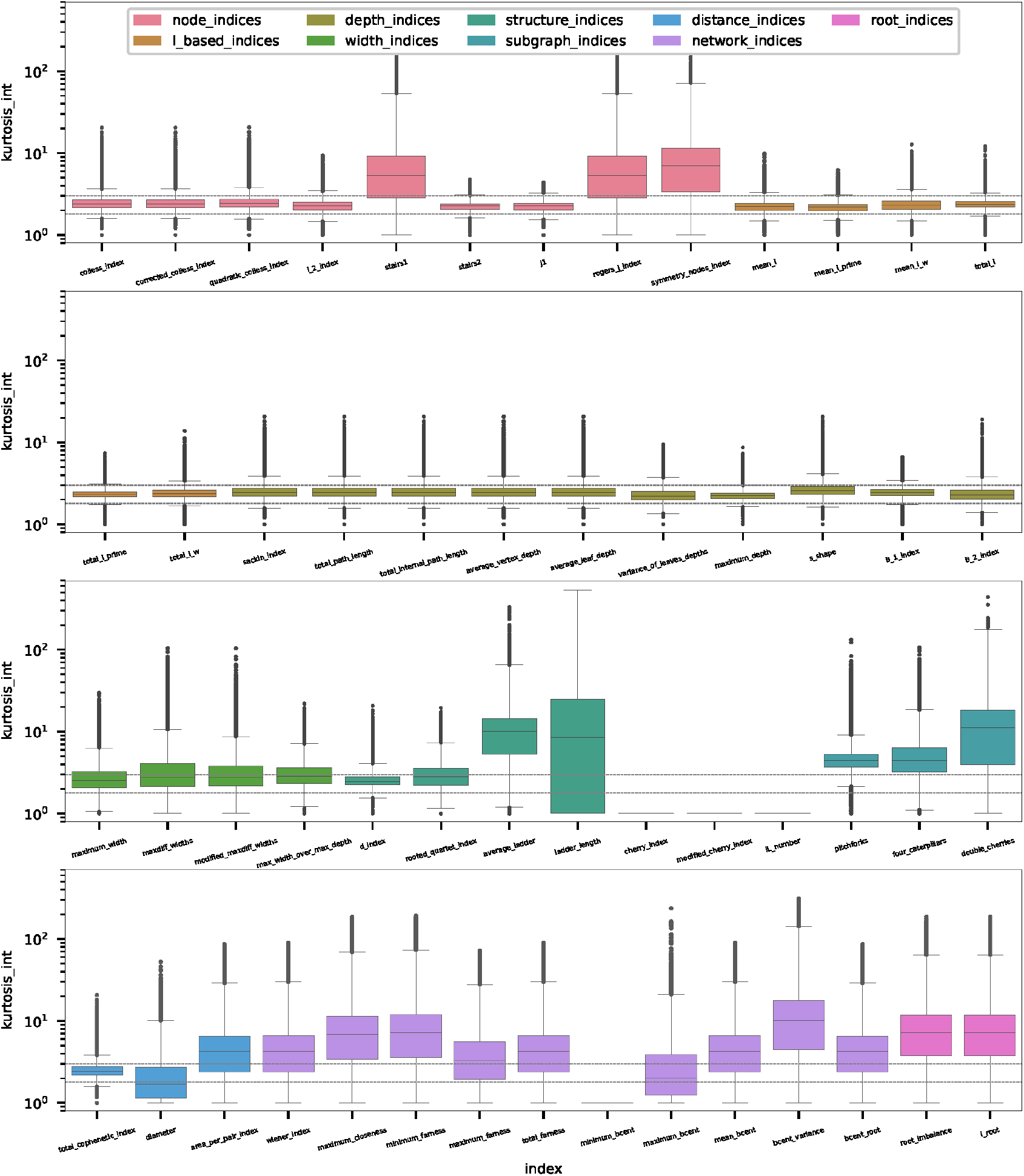
Kurtosis distributions for all tree shape indices over the trees under study considering *internal* roots only. The plots are grouped and colored according to the underlying approach of the indices (see Section 4.1). A dashed line at level 3 indicates the kurtosis of the normal distribution. The y-axis is logarithmically scaled. A horizontal line at level 10^0^ = 1 indicates, that the index behavior is entirely stable. That is, the same value is obtained if the root is placed on any internal branch.

**Fig. 17:**
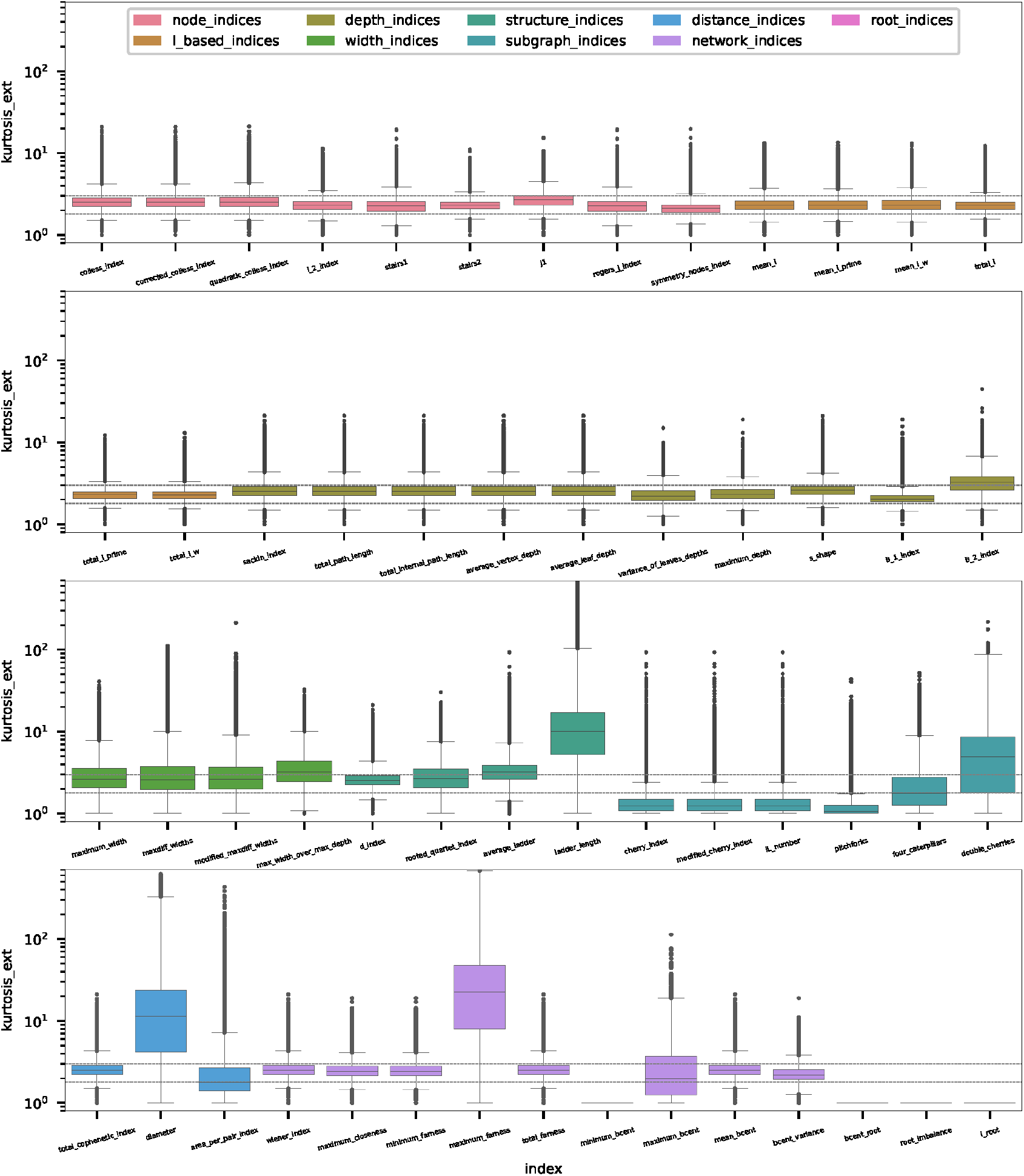
Kurtosis distributions for all tree shape indices over the trees under study considering *external* roots only. The plots are grouped and colored according to the underlying approach of the indices (see Section 4.1). A dashed line at level 3 indicates the kurtosis of the normal distribution. The y-axis is logarithmically scaled. A horizontal line at level 10^0^ = 1 indicates, that the index behavior is entirely stable. That is, the same value is obtained if the root is placed on any external branch.

**Fig. 18:**
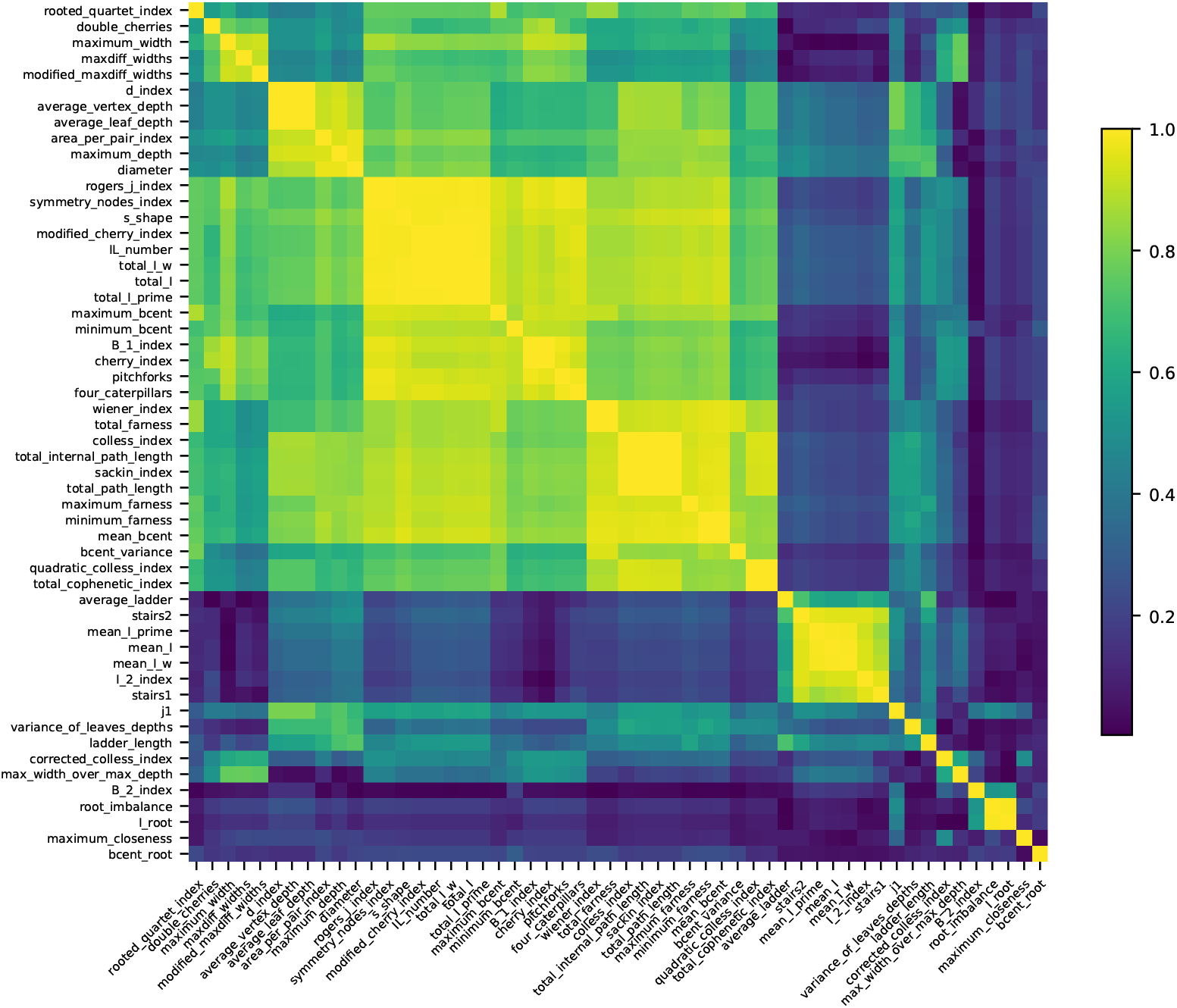
Pairwise Pearson correlations of tree shape indices. Indices with high correlations are grouped together. The correlations are obtained using all rooted trees under study.

**Fig. 19:**
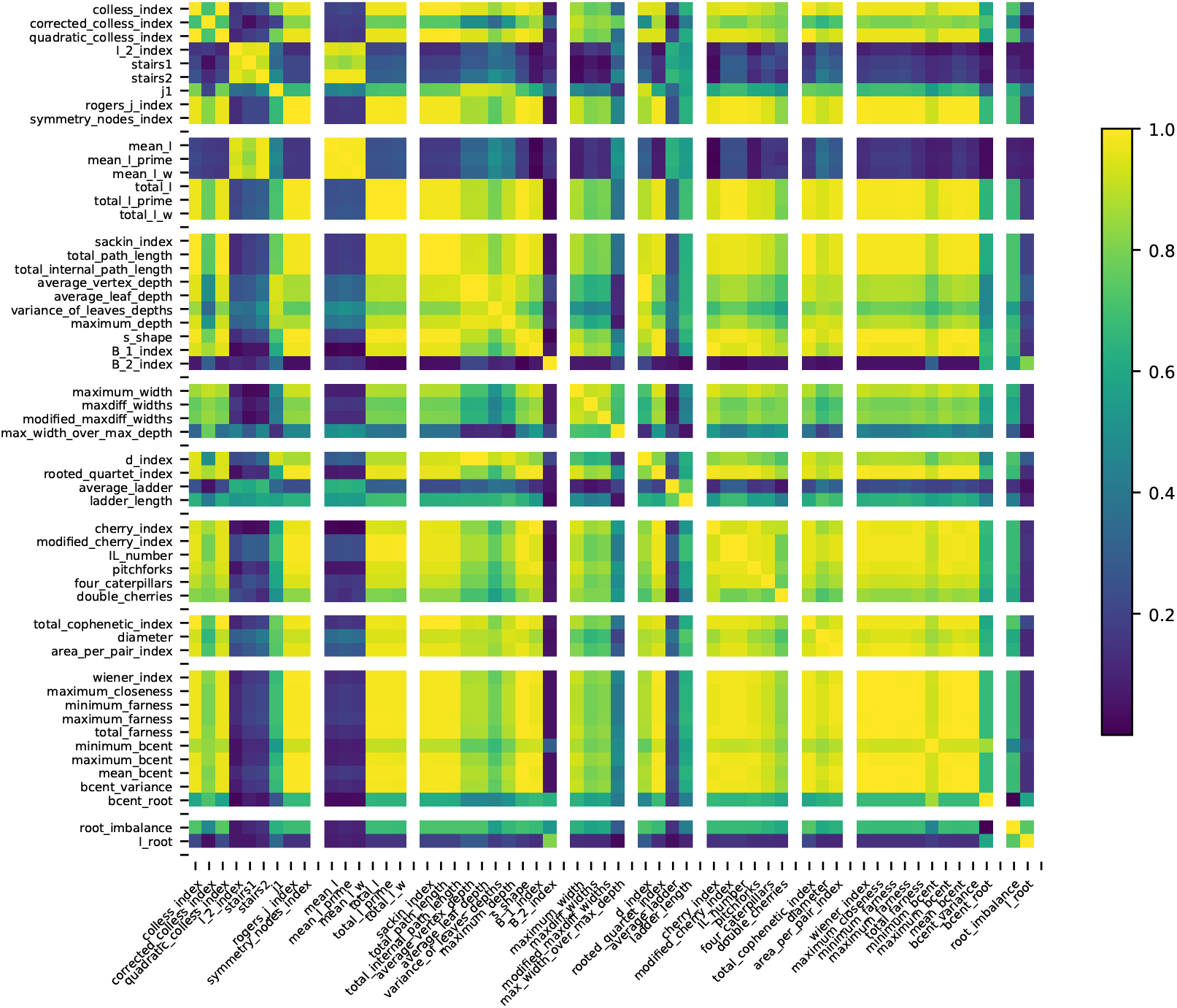
Pairwise Spearman Rank correlations of tree shape indices. The indices are grouped according their conceptual approach (see Section 4.1). The correlations are obtained using all rooted trees under study.

**Fig. 20:**
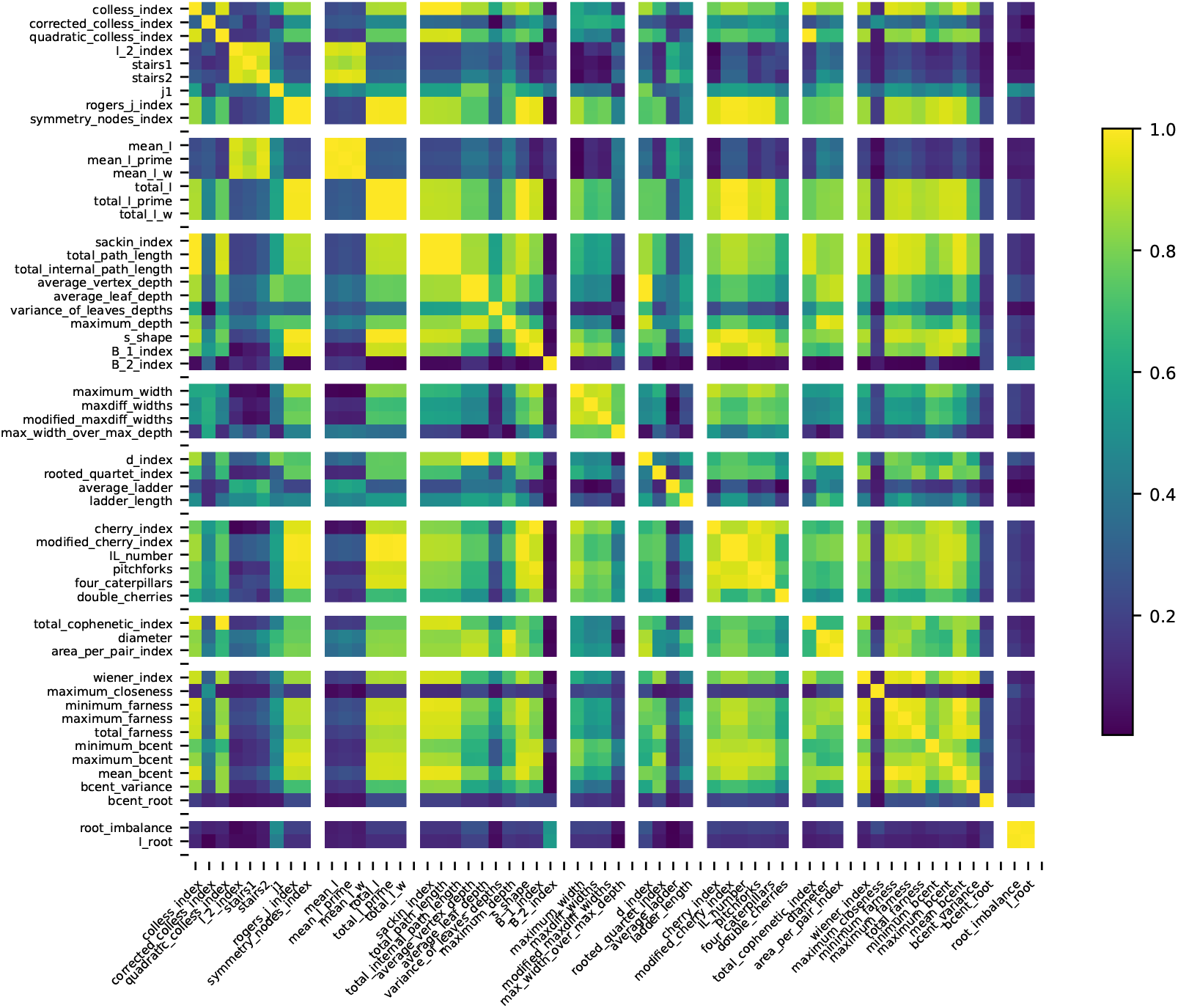
Pairwise Pearson correlations of tree shape indices. The indices are grouped according their conceptual approach (see Section 4.1). The correlations are obtained using all rooted trees under study.

**Fig. 21:**
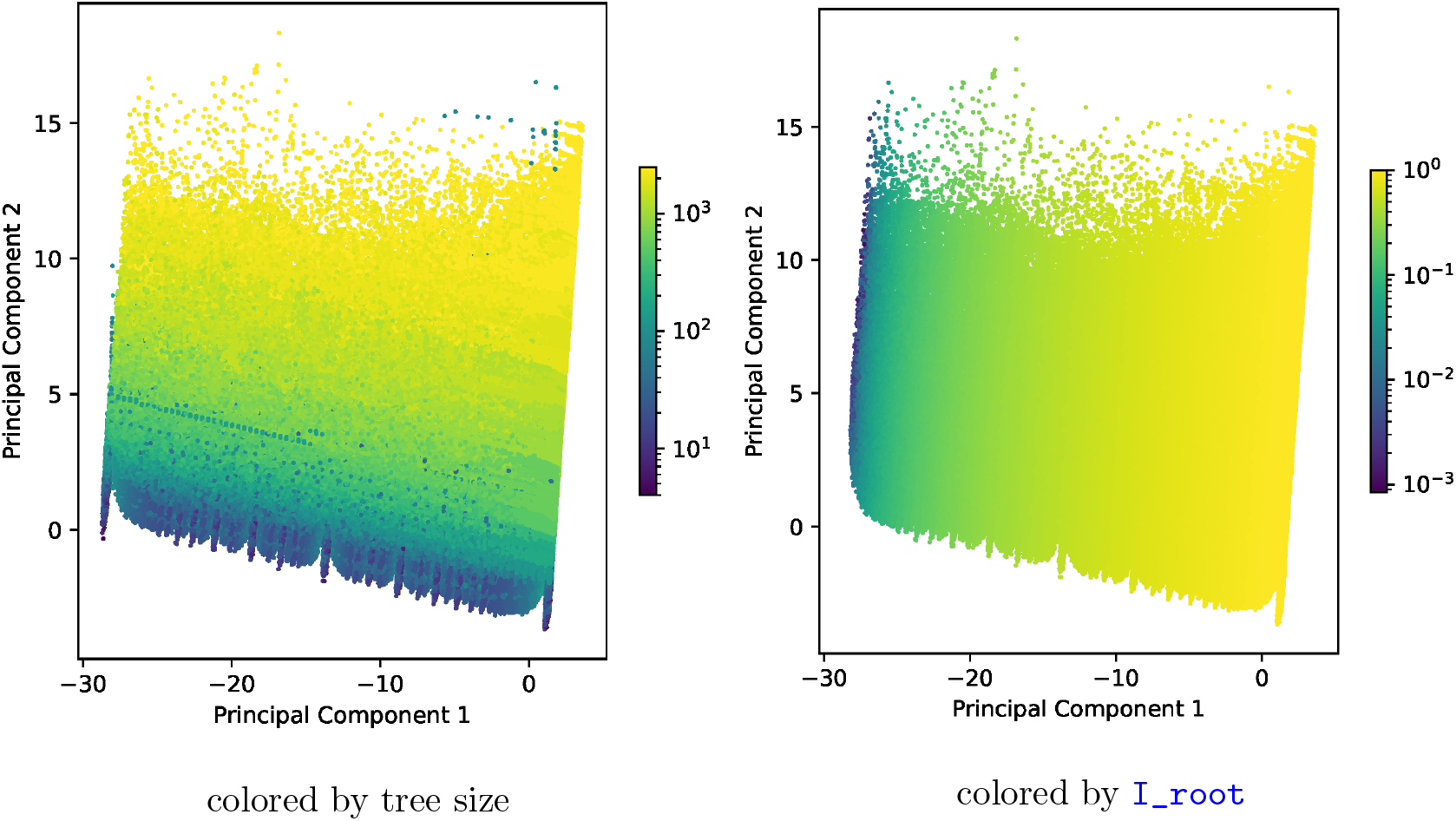
Results of our PCA. In each plot, there is one marker for each unrooted tree. The axes correspond to the principal components. The markers are colored according to the values of tree size and `I_root`.

In our final `treeshapy` implementation, the aforementioned node properties are not calculated via a pre-processing step, but on-demand at the first time they are required. We further exploit the dependencies of some indices not only on node properties but also on other indices (see Table 7). The `stairs1` index, for example, can by obtained as `rogers_j_index` divided by (*n* − 1). In practice, these dependencies induce additional speedups that are not reflected by our benchmark above.

A comparison to the runtimes of the `treestats` R library is provided in Fig. 15. The `treestats` library is based on a highly optimized C++ kernel and thus substantially outperforms `treeshapy` in terms of runtime.

Yet, we regard `treeshapy` as a useful complementary tool. As our library is written in Python, it extends the comprehensive Python code base for phylogenetics [36, 14, 58, 3]. Furthermore, the availability of two tools that offer analogous functions, but were independently developed allows for cross-validation and -verification (see Appendix A.2) that will allow to improve both tools. Although `treeshapy` runtimes are higher than those of `treestats`, they are still feasible for most application scenarios. For instance, calculating all indices on a single rooted tree with 100 leaves requires less than 15 milliseconds on average in our benchmark.

### A.2 Verification

To verify that `treeshapy` correctly computes all tree shape indices, we apply two different approaches: First, we use six different topologies, each with six tips (taken from Fischer et al. [24]). For each of these topologies, we manually calculate all tree shape indices and check whether we obtain the same values using `treeshapy`.

In addition, we use the R library `treestats` to evaluate the indices that are implemented in both `treeshapy` and `treestats` for all rooted trees under study. We ensure that the results from the two libraries differ by no more than 0.001. Note that we omit the `s_shape` since the libraries differ in an implementation detail (base of the logarithm). Further we cannot assess the correctness of the `colijn_plazotta_rank` due to large integer overflows.

The empirical trees under study are all fully bifurcating. However, `treeshapy` also allows to evaluate some indices on multifurcating trees. To also verify the correctness of this option, we sample 200 rooted tree topologies with multifurcations under the Yule model. For these trees, we evaluate all indices that can also be calculated on multifurcating trees. We compare the results obtained with `treeshapy` andţ`reestats` as described above.

### A.3 Usage Example

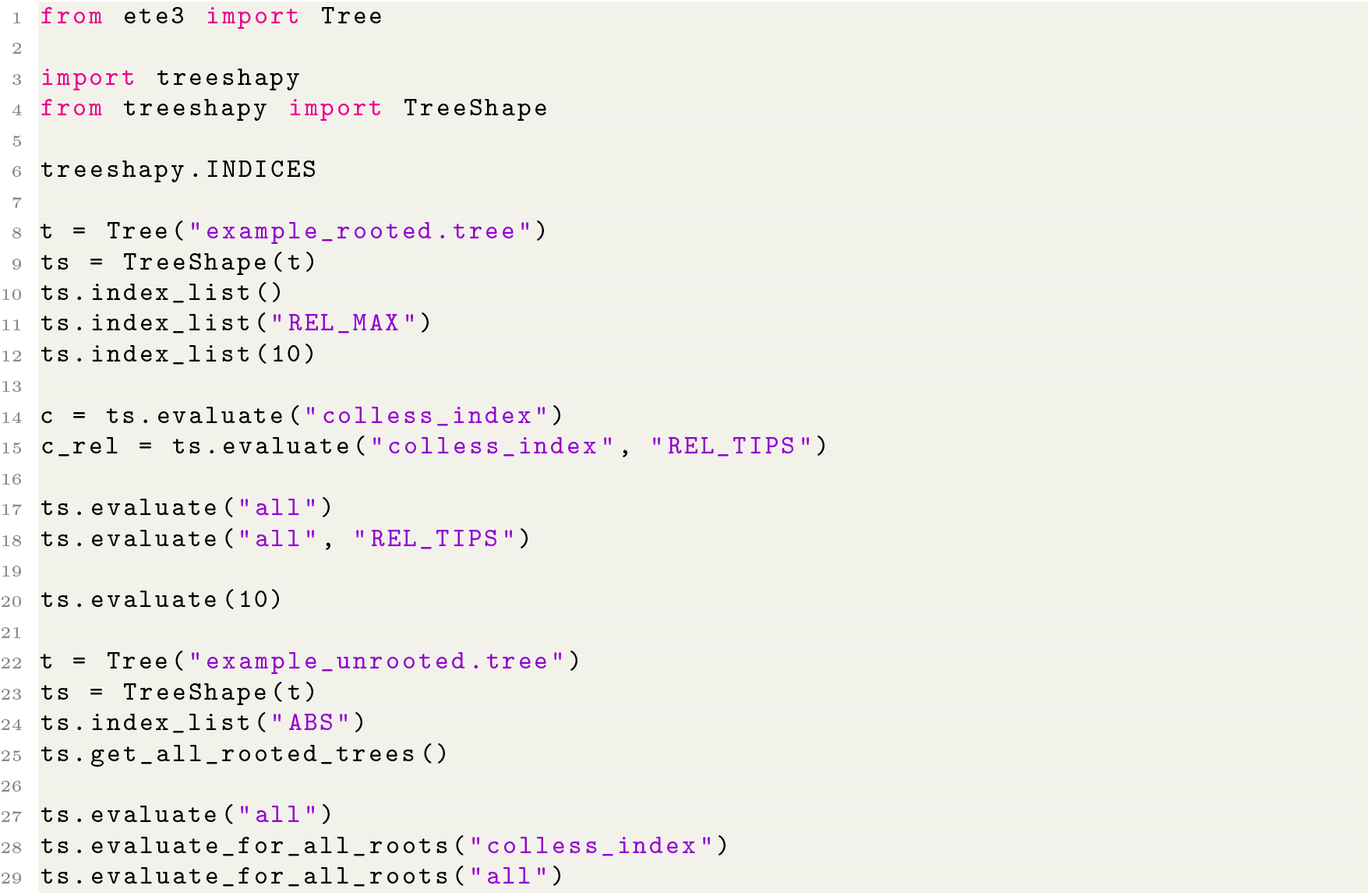

## B Index Properties

## C Applications of Tree Shape Indices

## D Additional Results

### D.1 Principal Component Analysis

The tree space is high dimensional, which yields comparing tree topologies particularly challenging. However, the plethora of tree shape indices allows to perceive the tree space in a different way. Instead of characterizing the trees by their discrete topologies, we regard each tree as a vector containing corresponding values for multiple tree shape indices. The resulting vector space is also high dimensional, but it allows to apply statistical methods for dimensionality reduction, such as *principle component analysis (PCA)*. Based on the pairwise index correlations (see Section 2.4), we determine a subset of 10 indices with minimal pairwise correlation (see Appendix D.1). For each rooted tree, we construct a vector which contains the corresponding values for these indices. On the resulting 10-dimensional space, we then conduct a PCA. The quality of a PCA can be measured by means of the **relative explained variance** [39]. For our PCA, we obtain a relative explained variance of 0.600 (0.196) for the first (second) principal component.

Next, we determine the correlation between the first two principal components and the tree shape indices. We find that, the second principal component strongly correlates with the tree size (Pearson correlation 0.956). This shows that most tree shape indices depend on the tree size, thus confirming our observations from Section 2.3. The first principal component yields a high correlation with `I_root` (Pearson correlation 0.997). `I_root` is the index among the 10 selected indices that correlates the least with tree size. Despite the high explained variance, the resulting principal components therefore do not provide a meaningful projection of the tree space into two-dimensional space. Instead, the PCA confirms that the vast majority of tree shape indices is highly correlated with tree size.

#### Subsets of Indices with Minimum Correlation

For *k* ∈ [2, 10], we determine index sets of size *k* with minimal pairwise correlations. Such a set minimizes the sum of the pairwise Spearman rank correlations among all index sets of size *k*. Note that these subsets exclude the ranking indices. These indices can yield extremely large integer values for which makes them infeasible for many practical applications.

*k* = 2: `mean_I_prime, cherry_index`

*k* = 3: `I_2_index, bcent_root, root_imbalance`

*k* = 4: `corrected_colless_index, stairs1, average_ladder, I_root`

*k* = 5: `corrected_colless_index, stairs1, maxdiff_widths, average_ladder, I_root`

*k* = 6: `stairs1, B_2_index, maxdiff_widths, modified_maxdiff_widths, average_ladder, I_root`

*k* = 7: `stairs1, variance_of_leaves_depths, B_2_index, maxdiff_widths, max_width_over_max_depth, average_ladder, I_root`

*k* = 8: `stairs1, mean_I_prime, B_2_index, maxdiff_widths, modified_maxdiff_widths, average_ladder, cherry_index, I_root`

*k* = 9: `corrected_colless_index, I_2_index, stairs1, variance_of_leaves_depths,B_2_index, maxdiff_widths, modified_maxdiff_widths, average_ladder, I_root`

*k* = 10: `stairs1, mean_I_prime, mean_I_w, B_1_index, B_2_index, maxdiff_widths, modified_maxdiff_widths, average_ladder, cherry_index, I_root`

